# EPISPOT: an epigenome-driven approach for detecting and interpreting hotspots in molecular QTL studies

**DOI:** 10.1101/2020.09.21.305789

**Authors:** Hélène Ruffieux, Benjamin P. Fairfax, Isar Nassiri, Elena Vigorito, Chris Wallace, Sylvia Richardson, Leonardo Bottolo

## Abstract

We present EPISPOT, a fully joint framework which exploits large panels of epigenetic annotations as variant-level information to enhance molecular quantitative trait locus (QTL) mapping. Thanks to a purpose-built Bayesian inferential algorithm, EPISPOT accommodates functional information for both *cis* and *trans* actions, including QTL *hotspot* effects. It effectively couples simultaneous QTL analysis of thousands of genetic variants and molecular traits, and hypothesis-free selection of biologically interpretable annotations which directly contribute to the QTL effects. This unified, epigenome-aided learning boosts statistical power and sheds light on the regulatory basis of the uncovered hits; EPISPOT therefore marks an essential step towards improving the challenging detection and functional interpretation of *trans*-acting genetic variants and hotspots. We illustrate the advantages of EPISPOT in simulations emulating real-data conditions and in a monocyte expression QTL study, which confirms known hotspots and finds other signals, as well as plausible mechanisms of action. In particular, by highlighting the role of monocyte DNase-I sensitivity sites from > 150 epigenetic annotations, we clarify the mediation effects and cell-type specificity of major hotspots close to the lysozyme gene. Our approach forgoes the daunting and underpowered task of one-annotation-at-a-time enrichment analyses for prioritising *cis* and *trans* QTL hits and is tailored to any transcriptomic, proteomic or metabolomic QTL problem. By enabling principled epigenome-driven QTL mapping transcriptome-wide, EPISPOT helps progress towards a better functional understanding of genetic regulation.

## Introduction

Molecular datasets and annotation databases are growing in size and in diversity. In particular, genetic data are now routinely collected along with gene, protein or metabolite level measurements and analysed in molecular quantitative trait locus (QTL) studies, with the aim of unravelling the regulatory mechanisms underlying common diseases. However, these studies present additional complexities compared to classical genome-wide association studies (GWAS). First, they entail a very different statistical paradigm: while GWAS consider a single or a few related clinical traits, molecular QTL studies typically involve hundreds or thousands of molecular traits, regressed on hundreds of thousands of genetic variants. Second, they need to accommodate two types of genetic control: a variant may affect molecular products of genes in its vicinity (*cis* action) or products of remote genes (*trans* action), where the latter mode of control is typically much weaker and, hence, harder to uncover than the former. In particular, *pleiotropic* or *hotspot* genetic variants may exert weak *trans* effects on many molecular traits.

The current mapping practice only partially embraces the features of QTL studies. Indeed, widely-used marginal screening approaches^1,2^ suffer from a large multiplicity burden and tend to lack statistical power as they do not exploit the regulation patterns shared by the molecular entities, whereas joint modelling approaches^3,4^ are often limited by the computational burden implied by the exploration of high-dimensional spaces of candidate variants and traits. To manage this tension between scalable inference and comprehensive joint modelling, we recently proposed a variational inference approach, called ATLASQTL^5^, which explicitly borrows information across thousands of molecular traits controlled by shared pathways, and offers a robust fully Bayesian parametrisation of hotspots; its increased sensitivity and that of earlier related models have been demonstrated in different molecular QTL studies^4–7^.

In complement to the actual mapping task, biologists increasingly try to capitalise on the wealth of available *epigenetic annotation sources* to infer the functional potential of genetic variants. The standard strategy uses epigenetic marks mostly for prioritisation of hits derived from marginal screening: it consists in looping through all the loci with statistically significant associations and, for each locus, inspecting a few marks to decide on “a most promising” functional candidate genetic variant among all those in linkage disequilibrium (LD). This approach presents the following disadvantages: first, publicly available databases nowadays contain several hundreds of epigenetic annotations. Preselecting just a few may involve omitting others that are relevant, which may bias the conclusions. Second, even if a comprehensive inspection were feasible, the degrees of relevance of the annotations may be very uneven and may depend on the conditions, cell types, tissues, and even genomic regions considered, so it is unclear how to weight each contribution. In response to this, a number of model-based approaches leveraging epigenetic annotations have been proposed over the past decade, whether for genome-wide association studies (e.g., iBMU^8^, bfGWAS^9^, FINDOR^10^) or fine mapping (e.g., PAINTOR^11^, RiVIERA^12^). Despite this extensive development, no existing method provides a solution to our problem, namely, modelling the functional enrichment of *trans*-QTLs and hotspots, a task which is substantially more complex and elusive than for the functional enrichment of *cis* QTLs or GWA signals for a series of related phenotypes. All available modelling tools are designed for genetic mapping with one^8–11^ or a few^12^ traits at a time, while *trans*-QTL and hotspot mapping requires considering thousands of traits simultaneously. Moreover, many approaches only accommodate small numbers of candidate annotations by computational or statistical stability constraints^8,9^, or take as input GWA summary statistics rather than individual-level data, thereby not benefiting from the added statistical power obtained from jointly modelling the latter, along with the functional information^10–12^.

Our work enables large-scale inference for *cis*- and *trans*-QTL regulation using whole panels of external epigenetic annotations and argues that the epigenome can serve both to increase statistical power for QTL mapping and to shed light on the biology underlying the uncovered genetic map in a systematic manner. Specifically, it couples a fully Bayesian QTL mapping strategy, in which all loci and molecular traits are analysed jointly, with a principled leveraging of epigenetic information by treating this information as complementary *predictor-level* data that may inform the probability of genetic variants to be involved in QTL associations. As successfully demonstrated in the context of genetic mapping with clinical traits, suitable use of epigenetic information can boost the detection of weak associations and help in discriminating genuine signals from spurious ones caused by LD or other confounding factors^13,14^.

Our modelling framework, called EPISPOT, directly infers the role of *sparse sets of annotations* — from *hundreds of candidate functional annotations* — in the activation of both *cis* and *trans* mechanisms affecting *hundreds to thousands of molecular traits*. Importantly, it combines this epigenome-driven feature with a flexible hotspot modelling feature inspired from our previous work^5^, thereby offering a unified toolkit to refine the detection of hotspots, aided by the epigenetic information at hand. The base version of EPISPOT assesses the action of the annotations uniformly for the full set of analysed transcripts. However, for cases where a sensible partition into subsets of co-expressed molecular traits (*modules*^15^*) is available, we also develop a module* version of EPISPOT, which accommodates module-specific epigenetic action by estimating the contribution of the epigenetic marks to the QTL associations in each module.

Our take is that fully joint modelling is paramount to borrow information across loci, epigenetic marks and molecular traits with complex dependences, but this requires careful algorithmic considerations to ensure scalable inference while retaining accuracy. EPISPOT implements an adaptive and parallel variational expectation-maximisation (VBEM) algorithm, augmented with a simulated annealing scheme which effectively explores the multimodal parameter spaces induced by highly-structured data. This optimisation routine is purposely tailored to the analysis of genetic data with strong LD blocks, for which the inclusion of the epigenetic data has the greatest impact. Our framework also constitutes a novel tool for *interpreting* (i) the detected *trans*-acting and *hotspot* variants based on their overlap with the selected epigenetic marks, and (ii) the molecular traits under genetic control in light of these marks. This additional purpose of EPISPOT is key given that elucidating the mechanisms of action of hotspots is often as challenging as mapping them in the first place. Indeed, there is accumulating evidence that most genetic variants acting in *trans* lie in intergenic regions^16–18^, where functional roles are difficult to decipher. Moreover, the massive *trans*-gene networks under genetic control are thought to be subject to subtle interplays, and researchers are often left with a variety of possible strategies to try to understand the interacting pathways between the genotype and underlying disease endpoints^19^. These strategies range from hypothesis-driven bottom-up approaches that start from isolated mechanisms and try to generalise them (e.g., based on *cis*-mediation hypotheses), to agnostic top-down approaches that directly model the whole system in view of teasing apart its fundamental components (e.g., based on graphical modelling approaches)^20^. Our approach provides an alternative anchor towards decoding the complex networks controlled by hotspots, namely via the epigenetic marks found to be informative for the genetic mapping.

EPISPOT is not targeted at genome-wide discovery but at effecting refined QTL mapping and hotspot prioritisation, based on genomic regions — hereafter called *candidate loci* — harbouring SNPs thought to be involved in QTL regulation. A crucial distinction with the existing enrichment approaches is that the candidate loci do not correspond to a previously-determined list of QTL hits but are *whole genomic regions*, which can involve hundreds of genetic variants (most of them with no QTL activity). EPISPOT exploits shared epigenetic signals across these regions to then select QTL hits with an increased statistical power.

Importantly, fruitful applications of EPISPOT, that can successfully decipher part of the molecular regulation machinery, require problems where the signal-to-noise and density of epigenetic/QTL signals are sufficient. In this work, we will describe extensive simulation experiments to highlight the benefits of using epigenetic information when available for a panel of regulation scenarios, and we will question the conditions under which inference is adequately powered to leverage this information. We will therefore formulate guidelines for practical use and provide a software implementation of EPISPOT along with documented code for the data-generation procedure used in the simulation experiments.

Another key component of the present paper concerns illustrating and exploiting the advantages of EPISPOT in real molecular QTL conditions. We will conduct and discuss the findings of a thorough monocyte expression QTL (eQTL) study leveraging a panel of annotations, including DNase-I sensitivity sites identified in different tissues and cell types, Ensembl gene annotations and chromatin state data from ENCODE. In particular, by pinpointing context-relevant marks in a hypothesis-free manner, EPISPOT will allow us to disentangle key mechanisms pertaining to the lysozyme pleiotropic activity of chromosome 12 — an activity which, although reported in several studies, is so far left unexplained in terms of its functional and mediation processes. Obtaining such evidence without EPISPOT would involve the daunting task of evaluating the enrichment of candidate eQTL hits in each individual epigenetic mark; this would also have no guarantee of success since one-at-a-time inspection strategies are deprived of the enhanced statistical power obtained with a unified joint epigenome/QTL mapping strategy.

## Material and Methods

### Two-level hierarchical regression model

We consider a Bayesian model linking three data sources (Figure 1A) with two levels of hierarchy. The bottom level parametrises the QTL effects and the top level parametrises the epigenetic modulations of the primary QTL effects.

**Figure 1:**
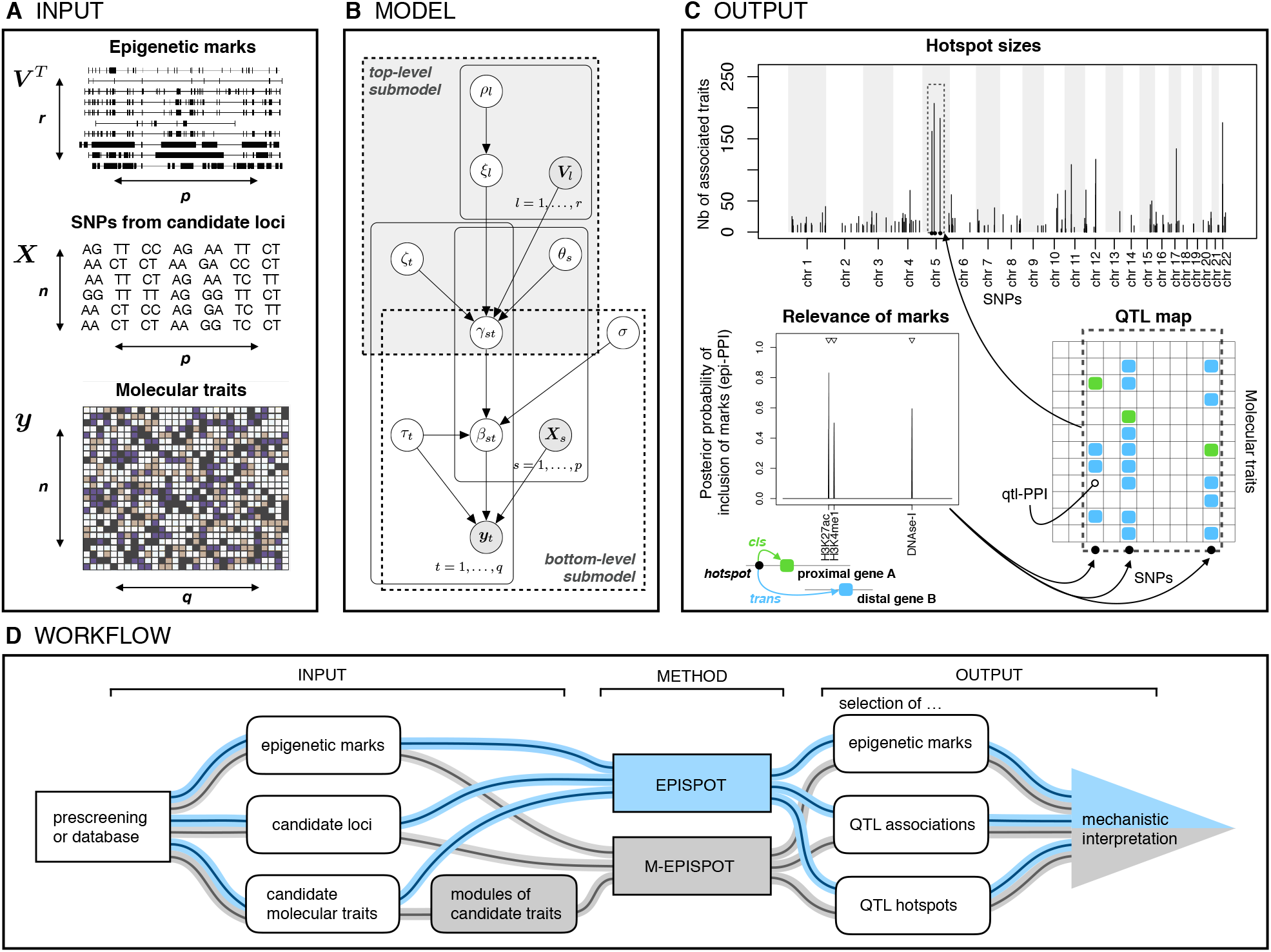
Overview of EPISPOT. A: Data input. Epigenetic annotations (predictor-level information) ***V***, genetic variants from candidate loci (candidate predictors) ***X***, molecular traits (responses) ***y***. B: Graphical representation for the two-level hierarchical model. The shaded nodes are observed, and the others are inferred. The top-level regression corresponds to the top plate; the probability of association is decoupled into a trait-specific contribution, *ζ*_*t*_, a SNP-specific contribution with a “hotspot propensity parameter” *θ*_*s*_ and an epigenome-specific contribution, *ξ*_*l*_, where ***V***_*l*_ is the vector gathering the observations of predictor-level epigenetic covariate *l* for all candidate SNP predictors ***X***_*s*_, *s* = 1, …, *p*. Parameter *β*_*st*_ models the effect between SNP ***X***_*s*_ and trait ***y***_*t*_, and *γ*_*st*_ and *ρ*_*l*_ are binary latent indicators for the QTL associations and epigenetic mark involvement, respectively. Parameter *σ* models the typical size of QTL effects and 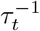 models the residual variability of trait ***y***_*t*_. C: Posterior output. Selection of epigenetic marks with a role in QTL regulation is carried out using the posterior probabilities of inclusion (epi-PPIs), pr(*ρ*_*l*_ = 1 | ***y***), *l* = 1, …, *r* (bottom left) and selection of associated SNP-trait pairs (aided by the marks) is carried out using the posterior probabilities of inclusion (qtl-PPIs), pr(*γ*_*st*_ = 1 | ***y***), *s* = 1, …, *p*; *t* = 1, …, *q* (bottom right). The hotspot Manhattan plot (top) reports the number of traits associated with each SNP (“hotspot size”), after using a selection threshold on the qtl-PPIs (e.g., FDR-based). D: EPISPOT workflow. Candidate loci and molecular traits are obtained from a preliminary screening or from existing databases, and supplied as input to the method along with epigenetic marks at the variants harboured by the loci. The algorithm is used with or without the module option depending on whether the traits are gathered into modules or not (M-EPISPOT in grey, resp. EPISPOT in blue). The output consists of sets of associated variants and traits, QTL hotspots and epigenetic marks relevant to the primary QTL associations, for given significance thresholds. It is then interpreted to generate mechanistic hypotheses about the functional processes underpinning the QTL associations.

Specifically, the bottom level hierarchy uses a series of conditionally independent spike-and-slab regressions to model the regulation of *q* molecular traits by *p* candidate genetic variants or single nucleotide polymorphisms (SNPs) for *n* samples:

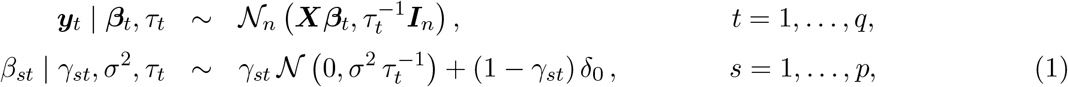

where ***y*** = (***y***_1_, …, ***y***_*q*_) is an *n* × *q* matrix of centred responses (molecular traits) and ***X*** = (***X***_1_, …, ***X***_*p*_) is an *n* × *p* matrix of centred candidate predictors for them (SNPs). Here, *δ*_0_ is the Dirac distribution and to each regression parameter *β*_*st*_ corresponds a binary latent parameter *γ*_*st*_ taking value 1 if and only if <u>S</u>NP *s* is associated with <u>t</u>rait *t*. Taking the posterior means of the latent parameters *γ*_*st*_ then yields marginal posterior probabilities of inclusion (qtl-PPIs, Figure 1C), pr(*γ*_*st*_ = 1 | ***y***), from which Bayesian false discovery rate (FDR) estimates can be obtained. Moreover, the precision parameters *τ*_*t*_ and *σ*^−2^ are assigned diffused Gamma priors.

The top-level hierarchy parametrises the effects of the epigenetic marks on the QTL probability of association via a second-stage probit regression on the probability of effects:

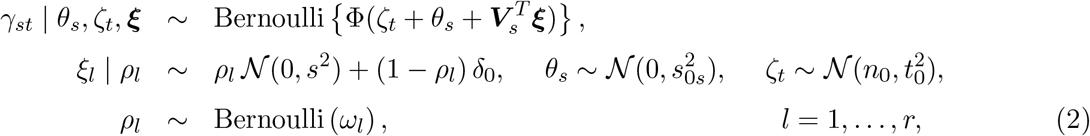

where Φ(·) is the standard normal cumulative distribution function and ***V*** = (***V***_1_,…, ***V***_*r*_) is a *p* × *r* matrix of (centred) predictor-level covariates (epigenetic marks).

Although prior information on the relevance of the marks for the QTL control can be accommodated if desirable, this is not required, as the use of a sparse prior on the mark effects ***ξ*** allows incorporating a large number of marks even though only a fraction may be responsible for genetic activity. In particular, if none of the marks are relevant, the QTL mapping will not suffer any bias from modelling the candidate marks (see simulation studies hereafter). Moreover, similarly as for the QTL effects, mark selection is easily achieved using posterior probabilities of inclusion, pr(*ρ*_*l*_ = 1 | ***y***), corresponding to the posterior means of the binary latent inclusion indicators *ρ*_*l*_(epi-PPIs, Figure 1C). This typically yields a sparse subset of marks, whose biological interpretation may help in understanding the mechanisms of action of the SNPs involved in the QTL associations.

### A parametrisation tailored to the detection of hotspots

In addition to embedding the predictor-level regression for the epigenetic effects, the top-level probit model in (2) also decouples the contributions of the predictors (SNPs) and the responses (molecular traits), namely, by involving a response-specific parameter, *ζ*_*t*_, which adapts to the sparsity level linked with each response ***y***_*t*_ and a predictor-specific parameter, *θ*_*s*_, which encodes modulations of the probability of association according to the overall effect of each predictor ***X***_*s*_. Parameter *θ*_*s*_ has a central role in pleiotropic molecular QTL settings as it represents the propensity of each predictor to be associated with multiple responses, i.e., its propensity to be a *hotspot*. Its Gaussian prior specification ensures closed-form updates, which is critical to the efficiency of the algorithm on large datasets. It also conveniently permits using a local-scale representation (via *s*_0*s*_) to prevent overshrinkage of large hotspot signals; see our previous work on the hierarchical modelling of hotspots, from which this formulation is borrowed^5^.

Here, the value of *s*_0*s*_ is set by empirical Bayes, and so are the epigenetic effect hyperparameters *ω*_*l*_ and *s*. The values of the hyperparameters *n*_0_ and *t*_0_ are chosen to induce sparsity, by specifying a prior expectation and a prior variance for the number of predictors associated with each response (Supplemental Material and Methods).

Hence, the EPISPOT model (1, 2) borrows information across the three types of entities (epigenetic marks, SNPs and molecular traits) in a unified manner, while providing interpretable posterior quantities, in particular qtl-PPIs and epi-PPIs, for the selection of each type of variables. It leverages the epigenome for two complementary purposes: (i) to enhance statistical power for QTL and hotspot mapping and (ii) to shed light on the biology underlying the genetic control, via the inspection of the selected marks.

### A modification for module-specific epigenetic contributions

The machinery of genetic control is complex and it is unlikely that the action of the epigenome on QTL regulation will uniformly affect the transcriptome. In particular, different groups of molecular traits may be governed by different functional mechanisms, involving different sets of epigenetic marks, to different degrees. When a partition into *modules* of genes (proteins or metabolites for pQTL or mQTL analyses, respectively) likely to be co-regulated is available to the analyst, it can be provided as input to the method which will then infer the annotation effects in a module-specific fashion, based on the following modification of the top-level model (2):

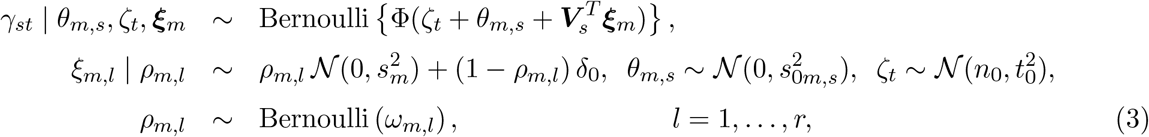

where *m* ∈*ℳ* is a module of traits, with *ℳ* a partition of *{*1, …, *q}* and *m 3 t*. Parameter ***ξ***_*m*_ then represents the epigenetic contribution of the *r* marks for the QTL associations involving the traits from module *m*. The hotspot parameter *θ*_*m,s*_ also accounts for the module structure: it represents the propensity of SNP *s* to be associated with few or many traits from module *m*. This encodes module-specific pleiotropic levels and also reflects the fact that a SNP controlling a given trait in a module is more likely to be also associated with related traits from the same module compared to traits outside the module.

The corresponding version of the algorithm — implementing model (1, 3) — is hereafter called M-EPISPOT when an explicit distinction with the base, module-free version — implementing model (1, 2) — is needed.

Different approaches, based on some prior state of knowledge, on specific optimisation methods or both, will typically yield complementary definitions of modules. In some instances, there will be obvious biological reasons backing up the obtained grouping, in others, no clear partitioning will emerge, in which case the analyst may choose to use the module-free version of the model. As there is no generic strategy for forming modules, it is important to understand the impact of such choices on inference. In particular, from a modelling point of view, a given module should ideally comprise co-regulated molecular traits, i.e., traits with shared genetic control, triggered by common epigenetic mechanisms. The top-level regression (3) will then represent the possible epigenetic effects underlying the functional mechanisms in the module, and module-specific epi-PPIs will be useful to select the marks involved in the regulation of each module. In particular, shared signals will be best leveraged when the molecular traits controlled by a given SNP belong to a same module. The simulation studies and the eQTL analysis will provide practical recommendations as well as analyses of sensitivity to module misspecification.

### A scalable purpose-built algorithm

The hierarchical model described above couples two levels of spike-and-slab regression, which accommodate three large spaces of SNPs, molecular traits and epigenetic marks, with possibly thousands of variables each. Careful algorithmic strategies are therefore critical to ensure that inference is accurate and scalable. To meet both requirements, we implement an adaptive variational expectation-maximisation (VBEM) algorithm and augment it with a simulated annealing procedure that efficiently explores the highly multimodal variable spaces formed by data with strong dependence structures.

VBEM algorithms were introduced by Blei et al. (2003)^21^ in the context of Dirichlet allocation modelling. In short, they iterate between optimising empirical Bayes estimates (in our case for the hotspot propensity and epigenetic effect hyperparameters) and running a variational algorithm for the remaining parameters, given the updated empirical Bayes estimates.

We present hereafter the algorithm in its general module-based form (M-EPISPOT); omitting the index *m* and taking *M* = 1 gives the base version with no module partitioning (EPISPOT).

Let ***v*** = (***β, τ***, ***γ***, *σ*^2^, ***θ, ζ, ξ, ρ***) denote the parameters for model (1, 3), and let ***η*** = (***η***_1_, …, ***η***_*M*_) denote the second-stage model hyperparameters, with 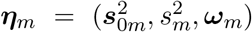 for module *m* = 1, …, *M*. We propose estimating ***η*** via an empirical Bayes procedure, by finding

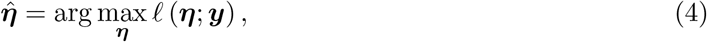

where *ℓ* (***η***; ***y***) = log *p* (***y*** | ***η***) is the marginal log-likelihood. Computing (4) analytically for our model would require high-dimensional integration and thus is infeasible. Our VBEM algorithm circumvents this by coupling the empirical Bayes estimation of the hyperparameter ***η*** with a variational inference scheme that simultaneously infers the model parameter vector ***v***. The procedure implements alternating optimisations of the variational lower bound

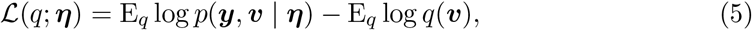

where *q*(***v***) is the variational density for 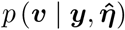 for a current estimate 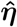 and E_*q*_ (*·*) is the expectation with respect to *q*(***v***). More precisely, it initialises the parameter and hyperparameter vectors ***v***^(0)^ and ***η***^(0)^, and alternates between the E-step,

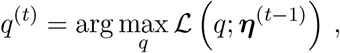

using the variational algorithm for obtaining *q*^(*t*)^ at iteration *t*, and the M-step,

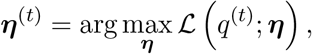

until convergence of ***η***^(*t*)^. In our case, the updates for the M-step are obtained analytically by setting to zero the first derivative of *ℒ (q*^(*t*)^; ***η***) with respect to each component of ***η***. This only requires computing and differentiating the joint likelihood term E_*q*_ log *p*(***y, v*** | ***η***) in (5), as the entropy term −E_*q*_ log *q*(***v***) is a function of ***η***^(*t*−1)^ and is constant with respect to ***η***.

Variational inference is typically orders of magnitude faster than classical Markov chain Monte Carlo inference^5,6,22^ for comparisons on GWA and molecular QTL models. Some computational cost is added for VBEM algorithms as each E-step requires running the variational algorithm until convergence. Moreover, the two regression levels of our model (1, 2) or (1, 3) necessitate the exploration of a very large parameter space, which is complex and time-consuming for any type of inference.

We consider two strategies to overcome this burden. First, we substantially reduce the runtime of the within-EM variational runs by using an adaptive stopping criterion, namely, starting with a large tolerance and dynamically decreasing it according to the convergence state of the overall EM algorithm. The second strategy applies to the module version of our algorithm: the specification in (3) suggests that its hyperparameters may be estimated reasonably well by restricting the VBEM scheme to subproblems corresponding to each module, i.e., applying model (1, 2) to the subsets of responses ***y***_*m*_ separately for obtaining the corresponding empirical Bayes estimates ***η***_*m*_, *m* = 1, …, *M*. In addition to accelerating hyperparameter estimation for each module (as the model is much smaller), this has the advantage of allowing parallelisation across modules. Once all module hyperparameters are estimated, they are inserted into model (1, HYPERLINK \l “bookmark3” 3) and variational inference is run on the entire dataset.

Strong posterior multimodality can be induced by dense genotyping panels with marked LD structures, whereby the inclusion of epigenetic information is particularly beneficial to disentangle the genetic contributions. To robustly infer signals from problems with strong data dependence structures, we augment all variational schemes with a simulated annealing routine^23,24^. Annealing introduces a so-called *temperature* parameter to index the variational distributions and control the level of separation between their modes, thereby easing the progression to the global optimum. In practice, we start with a temperature *T*_0_ to flatten the posterior distribution and sweep most local modes away, and we then lower it at each iteration, until the original multimodal distribution, called the *cold* distribution, is reached. Finally, to ensure stable inference, our routine excludes redundant SNPs and marks (i.e., displaying perfect collinearity with other SNPs/marks) prior to the run. Moreover, constant marks or marks which concern less than a given proportion of SNPs (default 5%) are also discarded before the analysis as insufficiently informative.

A sketch of the algorithm and the full derivation of the annealed VBEM updates are in the Supplemental Material and Methods. The algorithm is implemented as a publicly available R package with C++ subroutines^25^. Both the EPISPOT and M-EPISPOT versions run within seconds to few hours depending on the numbers of loci, molecular traits and epigenetic marks.

### Recommended use

EPISPOT is a refining tool for the detection and interpretation of QTL and hotspot effects. It is meant to be used for joint analysis of preselected genomic regions (*candidate loci*) and transcripts believed to be under genetic control (Figure 1D). Different approaches can be considered to obtain loci of interest. Public databases can be employed to form loci of given size around previously identified hits, provided this information is available for the condition, tissue or cell type at hand. An alternative approach is based on a preliminary application of ATLASQTL^5^ or another screening method, ideally on an independent dataset. If no independent dataset is available to the analyst, useful research hypotheses may still be obtained by running the prescreening step on the same dataset, prior to running EPISPOT. However, results should then be considered as exploratory, since this procedure interrogates the same data twice, which is subject to overfitting.

## Results

### Data generation and simulation set up

The series of simulation studies presented in the next sections have the dual purpose of (i) illustrating the effectiveness of EPISPOT in learning from the epigenome when the epigenetic annotations at hand are sufficiently informative (first simulation study), and (ii) evaluating the method in weakly informative scenarios (second simulation study) or scenarios where the module partition supplied to M-EPISPOT is misspecified (third simulation study).

We simulate data so as to best emulate molecular QTL regulation and the role of the epigenome in triggering this regulation; the general data-generation procedure is detailed in the Supplemental Material and Methods and we further tailor it to each simulation experiment in their dedicated sections.

We use the following terminology when referring to the simulated association patterns:

- an *active SNP* is a SNP with at least one association with a molecular trait;
- an *active locus* is a locus which involves at least one active SNP;
- an *active trait* is a trait with at least one association with a SNP;
- an *active module* is a module which contains at least one trait involved in QTL associations;
- an *active mark* is a mark which triggers at least one SNP-trait QTL association;
- the *hotspot size* is the number of traits associated with a given hotspot SNP.

We benchmark our approach against two representative state-of-the-art methods for QTL mapping, namely, the fully joint Bayesian QTL method ATLASQTL^5^, which is also tailored to the modelling of hotspots but does not accommodate the epigenetic marks, and the widely-used marginal screening approach MATRIXEQTL^2^, which tests each SNP-trait pair one-by-one and does not involve any epigenetic information.

### A first illustration

We first describe the type of posterior output produced by EPISPOT and its performance in a simple problem where no modules are involved, i.e., the active epigenetic marks exert their influence on all associated SNP-trait pairs.

We simulate 32 datasets with an average of 600 molecular traits, *r* = 500 candidate epigenetic marks and 60 candidate loci, each comprising an average of 20 real SNPs for 413 subjects. A subset of 100 SNPs are active (between 0 and 3 per locus) and their QTL effects are triggered by *r*_0_ = 3 active marks. This is a strong assumption, which permits a direct illustration of our algorithm in a simple setting, but since it may be unrealistic, we will only use it as a starting point for the more complex numerical experiments that follow. To help interpretability in the context of the simulations, we also generate marks with positive effects only, i.e., *inducing* QTL activity and not repressing it (Supplemental Material and Methods). The QTL signals are relatively weak: for any given trait, the cumulated QTL effects are responsible for at most 25% of its total variance. Many active SNPs are hotspots; across all 32 replicates, the active SNPs are associated with a number of traits ranging from 1 (isolated QTL association) to 96 (large hotspot), with an average of 27 active traits per active SNP.

Figure 2 shows that EPISPOT could clearly discriminate the three active marks contributing to the QTL associations from the remaining *r* − *r*_0_ = 497 inactive marks. The partial receiver operating characteristic (ROC) curves also show that it outperforms ATLASQTL in terms of selecting associated SNP-trait pairs and hotspots. It is unsurprising given that ATLASQTL does not use any predictor-level information, yet it nevertheless confirms that EPISPOT can effectively exploit the marks to enhance the estimation of the primary QTL associations. MATRIXEQTL performs poorly compared to the two joint approaches EPISPOT and ATLASQTL, which is expected since, by design, it does not exploit the shared association signals across traits.

**Figure 2:**
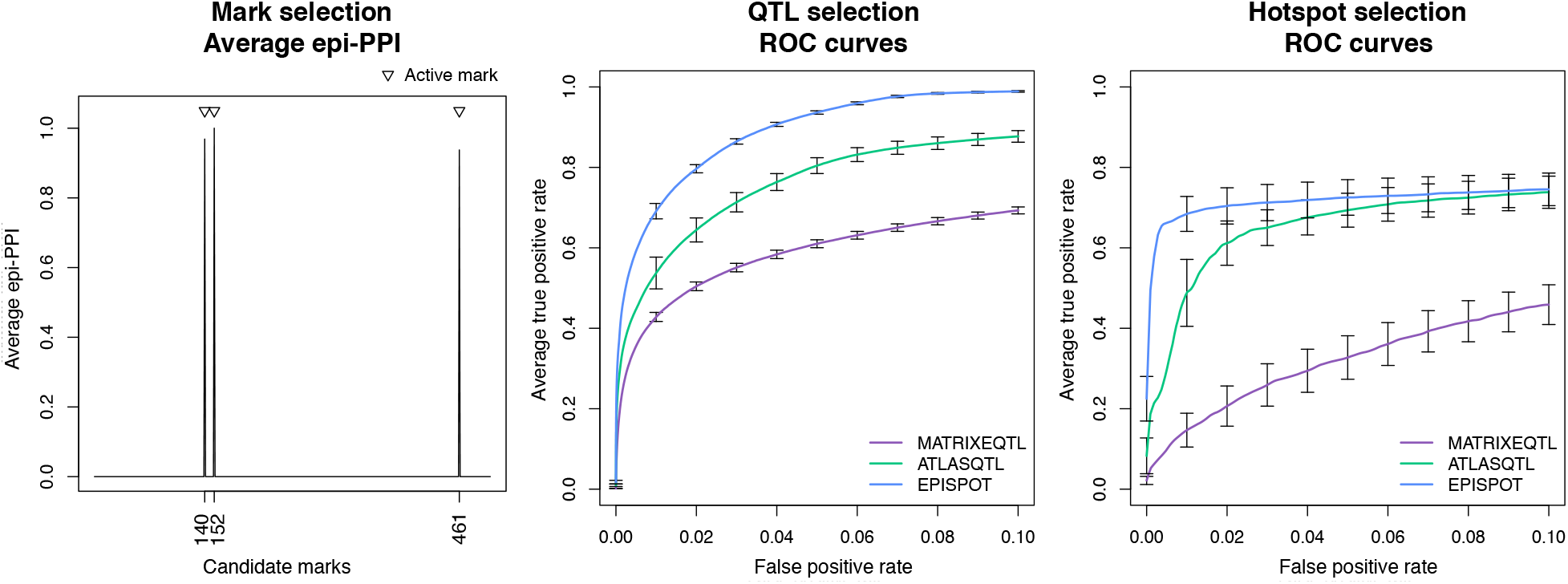
Performance for selection of epigenetic marks, pairs of associated SNPs and traits, and hotspots. Left: epi-PPIs for the marks averaged over 32 replicates. The three marks simulated as active are indicated by the triangles. Middle: average partial ROC curves for SNP-trait selection with 95% confidence intervals obtained from 32 replicates. EPISPOT is compared to the joint hotspot-QTL mapping method, ATLASQTL^5^, and the univariate screening method, MATRIXEQTL^2^, none of which makes use of the epigenetic marks. Right: idem for the selection of active SNPs (here, mainly hotspots).

We checked that EPISPOT and ATLASQTL display similar performance under simulation scenarios with no active mark: their 95% confidence intervals for the standardised partial area under the curve (pAUC) overlap, i.e., (0.74, 0.78) and (0.76, 0.79) for ATLASQTL, resp. EPISPOT (Supplemental Material and Methods). This further supports the observation that the improvement of EPISPOT seen in Figure 2 is attributable to an effective use of the three informative marks and not to other intrinsic differences between the two models; more evidence on this is provided in the next simulation experiment.

### Performance under varying degrees of epigenome involvement

Effectiveness in QTL mapping is subject to a number of interdependent factors pertaining to (i) the sparsity of the studied QTL network and magnitude of the QTL effects (ii) the amount of information contained in the data at hand (iii) the ability of the statistical approach to interrogate the data, i.e., by both leveraging and being robust to the dependence structures within and across genetic variants and molecular traits. When it comes to exploiting the epigenome to enhance statistical power, an additional level of complexity is introduced for determining the impact of the above factors on the analysis, and new questions arise as to whether the signal present in the data is sufficient to inform inference on the location of the relevant epigenetic marks and of the QTL associations potentially triggered by these marks.

In the previous simulation experiment, we generated data under the simplifying assumption that all QTL associations were induced by the epigenome, and to a degree to which the relevant marks would be detectable, as evidenced by the high epi-PPIs for the active marks and the power gained from leveraging this signal (Figure 2). Here, we focus on evaluating how the level of involvement of the epigenome in QTL activity impacts the detection of QTL effects and of the marks responsible for these effects.

We consider a series of QTL problems, each generated by replicates of 32, for a grid of response numbers and degrees of involvement of the epigenome in activating QTL control. More precisely, we simulate data with a number of traits sampled from a Poisson distribution with mean *λ* = 200, 400, 600, 800, 1000 or 1600, respectively, and 60 loci with 20 SNPs each and involving 100 active SNPs in total. We vary the proportion of active SNPs whose activity is triggered by epigenetic marks from *p*_epi_ = 0 (all QTL associations simulated independently of the action of the epigenome) to *p*_epi_ = 1 (all QTL associations simulated as the result of the action of the epigenome); see the Supplemental Material and Methods for the data-generation details. The typical pleiotropic pattern simulated is displayed in Figure 3 for the different choices of *p*_epi_ and problems with an average of *λ* = 600 traits.

**Figure 3:**
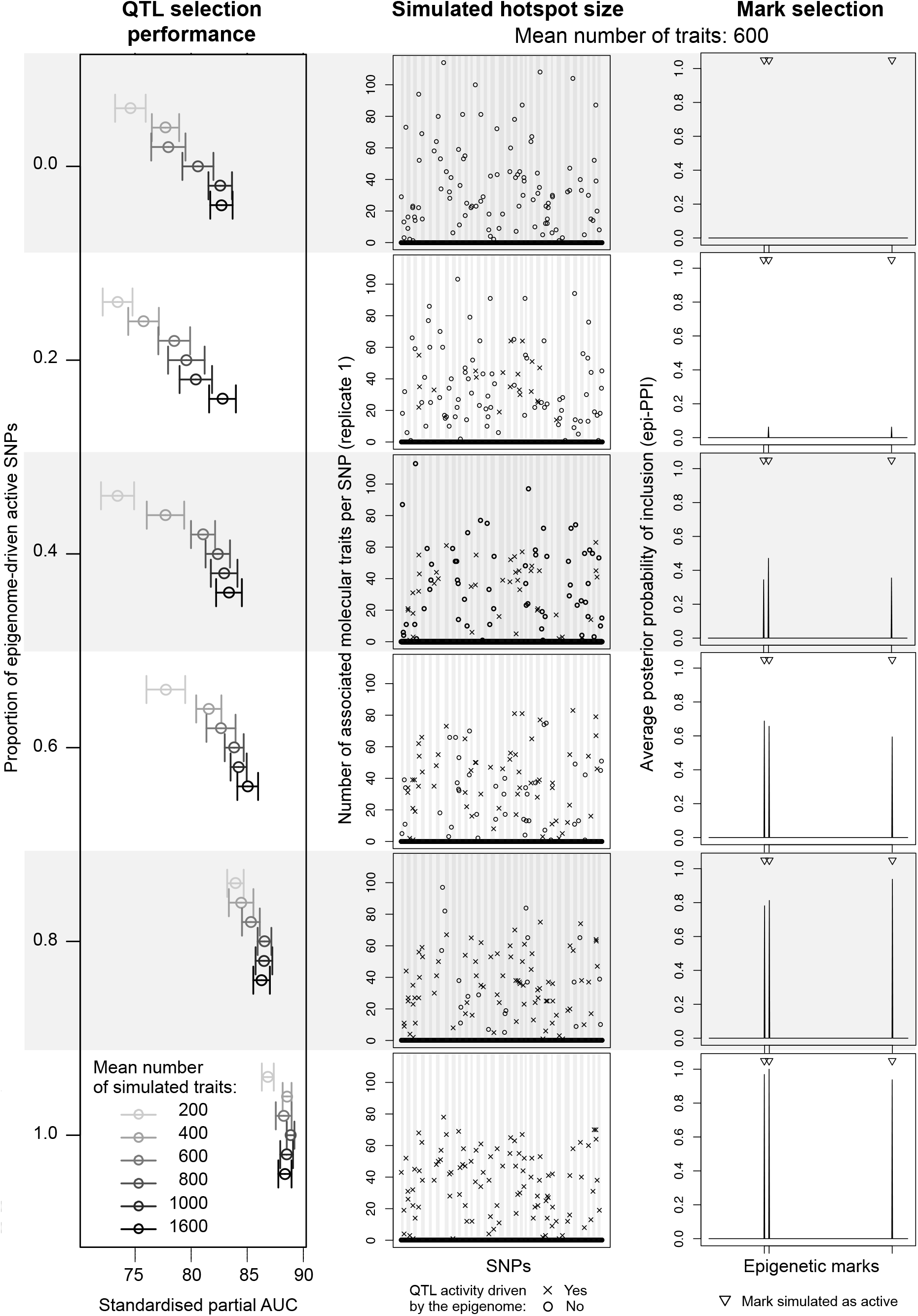
Performance of EPISPOT for a grid of numbers of traits and proportions *p*_epi_ of epigenome-driven active SNPs. Left: standardised pAUCs for the QTL selection performance with 95% confidence intervals. Middle: simulated hotspot QTL pattern for problems with an average of 600 traits (first replicate for each value of *p*_epi_). The crosses indicate hotspots whose activity is triggered by the epigenome and the circles indicate hotspots whose activity is independent of the epigenome. Right: average epi-PPIs, as inferred by EPISPOT for the simulated scenarios with an average of 600 traits.

Figure 3 also shows the performance for the selection of QTL effects in terms of standardised pAUC. It provides two separate layers of information: first, it illustrates again how EPISPOT is able to leverage the epigenetic marks to improve QTL mapping, and more so when the number of active SNPs triggered by these marks increases (top to bottom rows) since EPISPOT is then able to effectively borrow information across the mark-activated SNPs. This underlines the need for the relevant epigenetic marks to be sufficiently represented at causal variants so that the analysed data are informative about their involvement. It is therefore advised to use a reasonably large number of loci thought to be active and dense SNP panels (e.g., imputed SNPs, see the eQTL case study section), so the active SNPs are more likely to be included. Second, it shows that the joint modelling of all traits permits exploiting shared signals across these traits, thereby also improving statistical power, as reflected by the increased pAUCs for problems with larger numbers of traits in Figure 3. This is particularly true in presence of co-regulated molecular traits, a special case of which is the regulation of these traits by a single hotspot.

Figure 3 also indicates that, when the epigenetic signal is moderate to large (*p*_epi_ = 0.4, 0.6 or 1), EPISPOT is able to pick the active epigenetic marks from a large number of candidate marks, while setting the epi-PPIs of the inactive marks to zero. However, when the signal is weak (*p*_epi_ = 0.2), the active marks are barely detected, as expected. Importantly, though, in the *null scenario* where the epigenome plays no role (*p*_epi_ = 0), modelling the *r* = 500 inactive marks does not deteriorate the performance (Supplemental Material and Methods).

### Inferring module-specific epigenetic action

The simulation experiments presented next focus on evaluating M-EPISPOT, i.e., the module version of the algorithm which models module-specific epigenetic effects. They illustrate how statistical power and interpretability are enhanced when the structure underlying epigenome-driven QTL associations is exploited. They also evaluate the robustness of inference when misspecified module partitions are supplied to M-EPISPOT. This is particularly important given the uncertainty that often surrounds the definition of modules, as reflected by fact that different co-expression inferential tools often produce different module specifications.

We start with a simple example involving 60 concatenated loci of average size 40 SNPs and two modules of 50 simulated traits each. In the first module *m*_1_, the traits are largely co-regulated by hotspots whose activity is imputable to the epigenome. In the second module *m*_2_, only few traits are involved in isolated QTL associations, with no implication of the epigenome. Figure 4A illustrates the corresponding simulated QTL pattern restricted to the active SNPs, for the first data replicate. We evaluate the performance of M-EPISPOT with the following settings:

**Figure 4:**
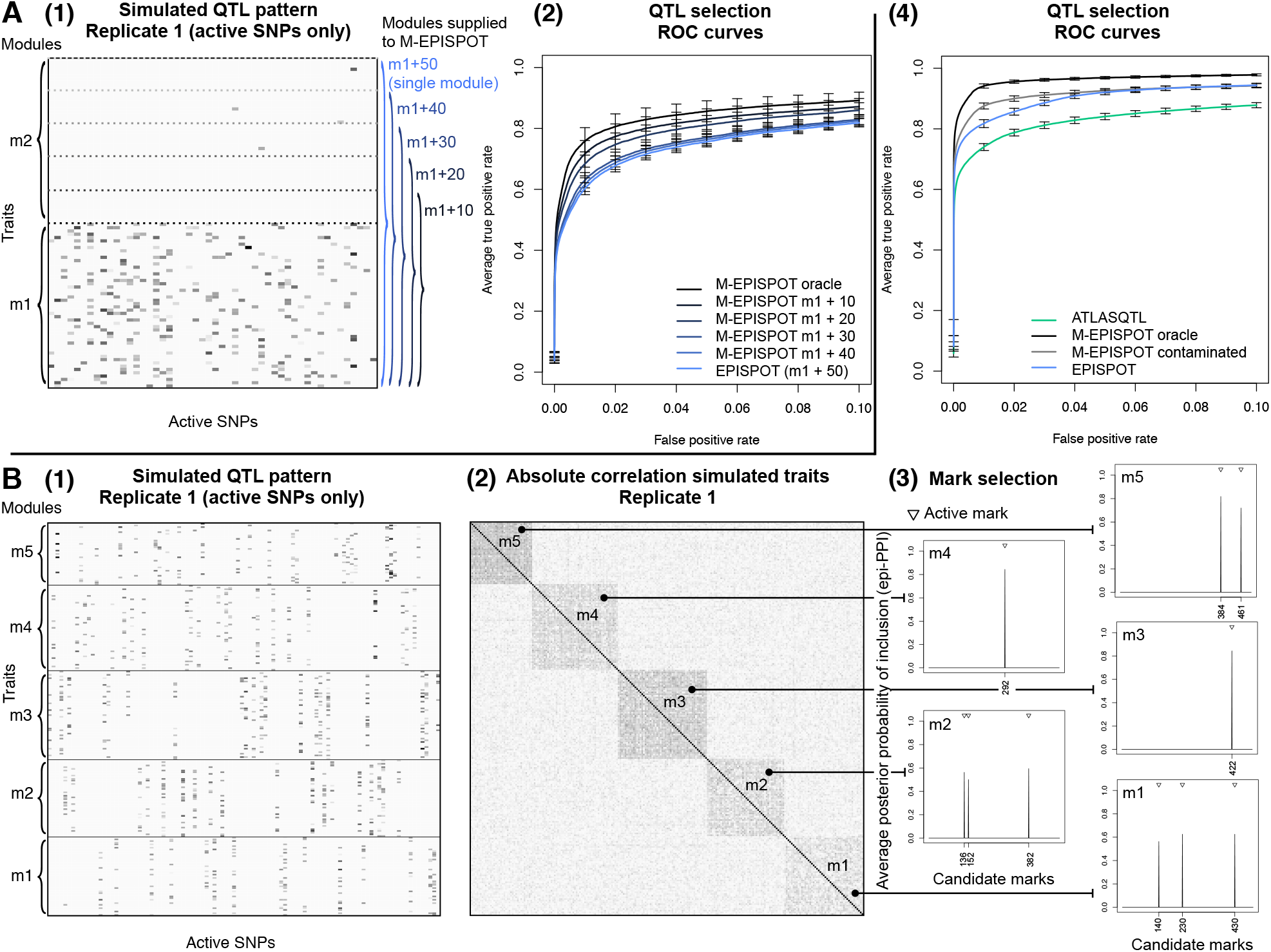
Performance of M-EPISPOT. A: Simulated scenario with two modules, whereby the first module *m*_1_ is contaminated by an increasing number of traits from the second module *m*_2_. Panel A(1) shows the simulated pleiotropic pattern for one replicate. The grey levels suggest the different QTL effect strengths of each active SNP (*x* axis) with the traits (*y* axis) from modules *m*_1_ and *m*_2_. The horizontal dotted lines mark the boundary between *m*_1_ and *m*_2_ for the misspecified module partitions supplied to M-EPISPOT. Panel A(2) shows the partial ROC curves (with 95% confidence intervals based on 32 replicates) for the QTL mapping performance obtained when supplying the different misspecified partitions shown in A(1) to M-EPISPOT. B: Simulation with five pleiotropic modules. Panel B(1) shows the simulated pattern for the active SNPs of one replicate. Panel B(2) panel shows the dependence structure of the simulated traits for one replicate. Panel B(3) shows the module-specific average epi-PPIs for the contribution of the epigenetic marks to the QTL effects. Panel B(4) shows the partial ROC curves for the QTL mapping, with 95% confidence intervals based on 32 replicates.

1. The oracle case, where we assume the simulated module partition *ℳ* = *{m*_1_,*m*_2_*}* to be known and provided it as input to M-EPISPOT;
2. the module-free case, where we perform inference with the base model EPISPOT which does not exploit the module partition;
3. a series of intermediate cases, where the module partition supplied to M-EPISPOT is misspecified, i.e., module *m*_1_ is contaminated with 10, 20, 30 or 40 traits from module *m*_2_ (Figure 4A). This mimics a real data scenario whereby the assignment of some traits to modules is difficult.

The ROC curves of Figure 4A show that leveraging information about the underlying module partition can improve significantly the detection of QTL effects. They also confirm the intuition that the impact of misspecified partitions on performance is a function of the degree of misspecification: for a given specificity, the power decreases smoothly with the number of inactive traits from module *m*_2_ contaminating module *m*_1_. From a modelling point of view, leaving all traits controlled by a same hotspot in a single module permits maximising the opportunities to learn the epigenetic contribution to the QTL activity by borrowing strength across co-regulated traits. It is advised to make use of prior information on pleiotropy when available in order to avoid splitting hotspot-controlled networks of traits into distinct modules.

The second simulation experiment considers a more general setting with five modules of average size 50. It compares ATLASQTL, EPISPOT, M-EPISPOT with the oracle module partition supplied and M-EPISPOT with a contaminated module partition supplied, i.e., where a fifth of the traits in the simulated modules are randomly re-assigned to the other modules.

Figure 4B leads to a conclusion similar to that of the previous example: the idealised scenario of the oracle module partition provided to M-EPISPOT yields the best performance, followed, in order, by the more realistic case of the contaminated partition, the EPISPOT run (with no module information) and finally, the ATLASQTL run which does not make use of any epigenetic information. Importantly, the fact that the module-free version EPISPOT outperforms ATLASQTL indicates that, even when the module structure is not employed, the method is still able to leverage the epigenome in order to improve the QTL mapping.

Figure 4B also shows how the marks responsible for the activation of the different modules are correctly recovered by M-EPISPOT. An inspection of these separate sets of marks provides a refined level of interpretability for a module-specific understanding of the genetic control. We will see in the eQTL analysis presented next how this can be particularly helpful to shed light on the mechanistic action of *trans* hotspots, when such hotspots are thought to control gene modules in a context-specific way.

### An epigenome-driven monocyte eQTL case study

In this section, we take advantage of EPISPOT in a targeted eQTL study to refine the detection and characterisation of genetic regulation in monocytes. Specifically, we analyse two independent datasets with transcript levels measured in CD14^+^ monocytes. Our study workflow is described in Figure 5A: we discover active loci in a *prescreening step* using the joint hotspot QTL mapping approach ATLASQTL^5^ in the first dataset (*n* = 413 samples^26^), and we then leverage the epigenome using EPISPOT in the second dataset (CEDAR cohort, *n* = 286 samples^27^) for an in-depth analysis of the genetic activity in the preselected loci.

**Figure 5:**
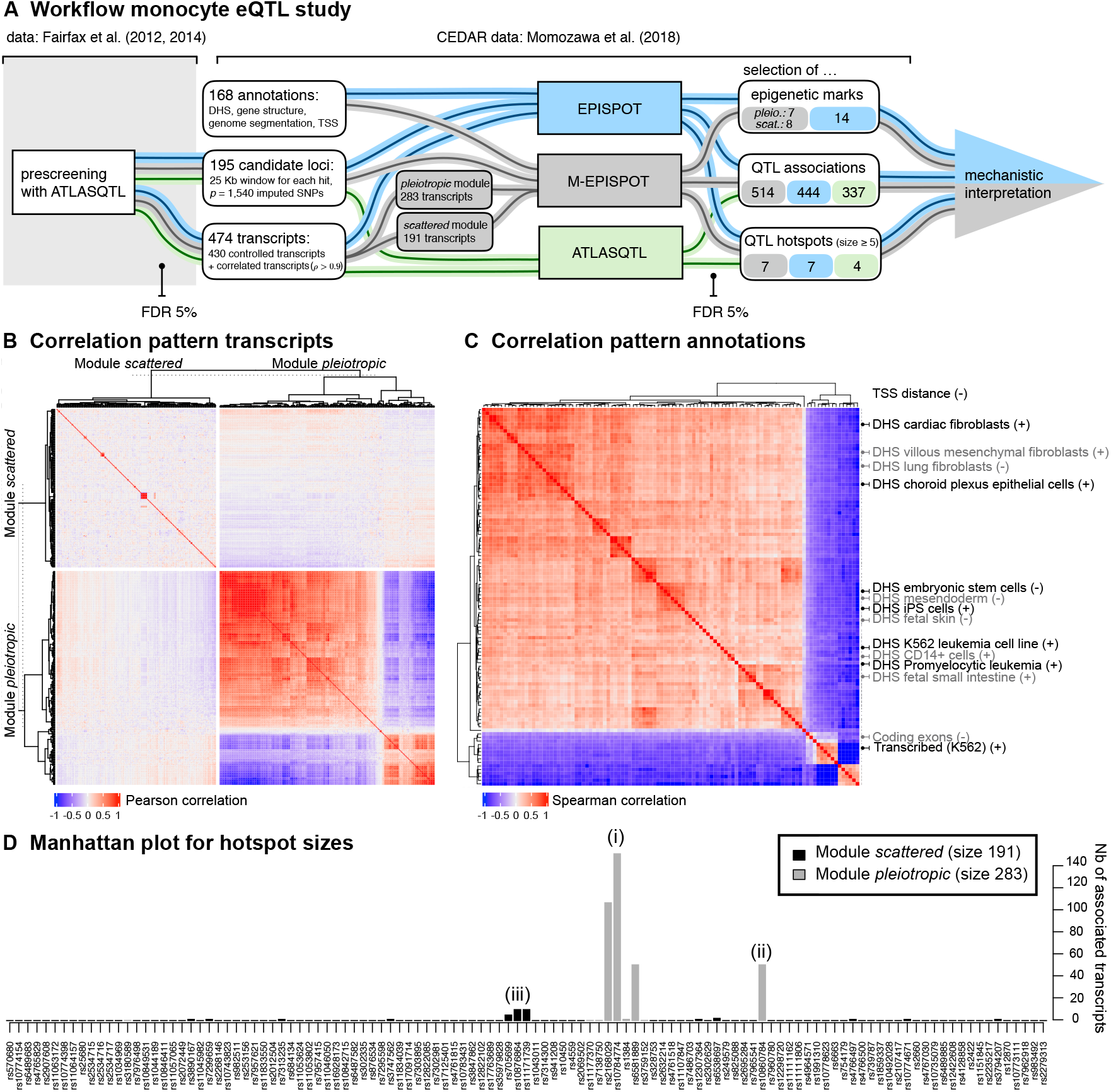
Overview of the monocyte eQTL case study. A: Workflow for the monocyte eQTL case study. Candidate loci from chromosome 12 and transcripts are obtained from a preliminary prescreening in the first dataset^26^ using the joint eQTL mapping approach ATLASQTL^5^ with a permutation-based Bayesian false discovery rate (FDR) of 5% for selecting pairs of associated SNP-transcript. The analysis is then performed in the second dataset (CEDAR)^27^. EPISPOT and M-EPISPOT select associated SNP-transcript pairs, QTL hotspots and epigenetic marks relevant to the primary QTL associations. This output is then interpreted as a whole to generate hypotheses about the mechanisms of action underlying these associations. B: Correlation of the analysed transcripts, according to their module membership. The *pleiotropic module* displays a strong dependence pattern, reflecting dense connections in the network controlled by the hotspots; the traits in the *scattered module* are mostly uncorrelated, which is unsurprising given that they are mainly controlled via isolated *cis* mechanisms. C: Correlation of the epigenetic annotations supplied to the method. All variables are binary, except the distance to the closest transcription start site (TSS) which is not included in the heatmap. Only the labels of the marks retained by M-EPISPOT are displayed; a heatmap with the full labels is provided in the Supplemental Material and Methods. The majority of the marks are DNase-I hypersensitivity sites (DHS) in different tissues and cell types. They tend to cluster together on the top-left 4*/*5 of the heatmap, and DHS in similar tissues/cell types also form subgroups. The remaining marks relate to gene structures and genome segmentation annotations. The labels indicated on the right are in grey and black depending on whether they were selected by M-EPISPOT as relevant for the *pleiotropic*, resp. *scattered module*. The (+) and (-) indicate positive, resp. negative effects of the marks, i.e., their triggering or repressive action on the primary QTL effects. Their relevance is discussed in the main text and in the Supplemental Material and Methods. D: Hotspot sizes (i.e., number of associated transcripts per SNP) as inferred by M-EPISPOT. Only the active SNPs (i.e., associated with ≥1 transcripts) are displayed. The grey and black colours indicate the module membership of the controlled transcripts. The numbers in parentheses refer to the discussion of the main text.

The epigenetic information consists of a panel of 168 annotation variables, compiling DNase-I sensitivity sites from different tissues and cell types, Ensembl gene annotations and chromatin state data from ENCODE. These variables display strong correlation structures within annotation types, as well as within tissues and cell types at a finer granularity level (Figure 5C). Details about the prescreening step, as well as the epigenetic, genetic and expression datasets are given in the Supplemental Material and Methods, and the eQTL associations for the prescreening and subsequent analyses are listed in Tables S1–S4.

In this case study, we concentrate our attention on the following key finding revealed by the prescreening step: chromosome 12 is highly pleiotropic, notably around the gene *LYZ*. This gene encodes lysozyme, a highly conserved enzyme with peptidoglycan-lytic activity that is robustly expressed in monocytes. The *LYZ* locus has already been reported as pleiotropic using several monocyte datasets^28–30^, but its functional role remains unclear. We will therefore exploit the epigenetic annotations within EPISPOT to shed light on the mechanisms of action of this locus as well as of other surrounding *cis*- and *trans*-acting loci.

#### The LYZ-region pleiotropy defines two modules of transcripts

A total of 977 eQTL associations, involving 350 unique SNPs on chromosome 12 and 430 unique transcripts genome-wide, were identified at FDR 5% from the ATLASQTL prescreening analysis of the first dataset. When mapped to the CEDAR dataset, the ATLASQTL eQTLs corresponded to 195 independent loci, comprising a total of *p* = 1, 540 SNPs (see Figure 5A and data-preparation details in the Supplemental Material and Methods). As highlighted in the second simulation study (Section “Performance under varying degrees of epigenome involvement”), supplying a dense panel of SNPs (here imputed SNPs) to EPISPOT is important to ensure a sufficient representation of the relevant epigenetic marks among the analysed SNPs.

We also mapped the prescreened transcripts to the CEDAR dataset. The *LYZ* -region pleiotropy defines two natural modules of transcripts, based on whether they are associated with SNPs in the vicinity of *LYZ* (< 1 Mb from it) or not, and further augmenting these modules with highly correlated transcripts (Supplemental Material and Methods). This module partition is driven by the following biological consideration: the peculiar pleiotropic QTL activity arising from the *LYZ* region may be triggered by specific epigenetic influences, which may differ from those triggering isolated (*scattered*) *cis* or *trans* effects outside the *LYZ* region; to reflect this, the modules are hereafter referred to as the *pleiotropic module* and the *scattered module*, respectively (Figure 5A). The correlation structure within and across the two modules supports this partitioning (Figure 5B). Namely, it indicates a strong co-expression of transcripts within the *pleiotropic module*, suggesting a dense network of genes whose connections may be attributed in large part to the shared QTL control exerted by the *LYZ* hotspots. Conversely, the transcripts in the *scattered module* display little co-expression, which is unsurprising given that they tend to be involved in isolated QTL effects (most transcripts are controlled by distinct genetic variants).

#### Overall comparison of methods and replication rates

We next refined our understanding of the eQTL structure in this region using the CEDAR dataset. To assess the sensitivity of inference to this module partition, we compared the results of the module-based algorithm, M-EPISPOT, with those of the base algorithm, EPISPOT, i.e., with no module provided as input. Moreover, to highlight the benefits of using epigenetic information, we also confronted these two runs with an ATLASQTL analysis of the same data. We employed the same settings for all three runs to set common grounds for comparison. In particular, we used a same permutation-based Bayesian FDR threshold of 5% for declaring QTL associations (Figure 5A and Supplemental Material and Methods). Importantly, the simulated annealing scheme implemented as part of the EPISPOT algorithm is specifically designed to handle the strong LD structures present in the dense SNP panel data and the block correlation structures among transcript levels (Figure 5B) and epigenetic marks (Figure 5C).

In the CEDAR dataset, the M-EPISPOT analysis of the two modules (*q* = 283 + 191 transcripts) and the 195 candidate loci (*p* = 1, 540 SNPs) identified 514 eQTL associations, involving a total of 267 unique transcripts and 82 unique loci (Table S2). In terms of independent replication of the prescreening hits, this corresponds to rates of 78.2% and 55.8% for the *cis* and *trans* QTL associations respectively. Using ATLASQTL instead of M-EPISPOT on the CEDAR data yielded 262 unique active transcripts and 80 unique active loci, with slightly lower *cis* and *trans* replication rates, namely 77.9% and 54.9% respectively (Table 1, Table S4). Similar observations were obtained for EPISPOT (Table 1, Table S3). Given the well-known difficulty to validate *trans* effects and the relatively small sample size of the CEDAR dataset (*n* = 289), these appreciable independent replication rates may result from the efficient joint modelling of all transcripts and SNPs achieved by M-EPISPOT, EPISPOT and ATLASQTL.

**Table 1:**
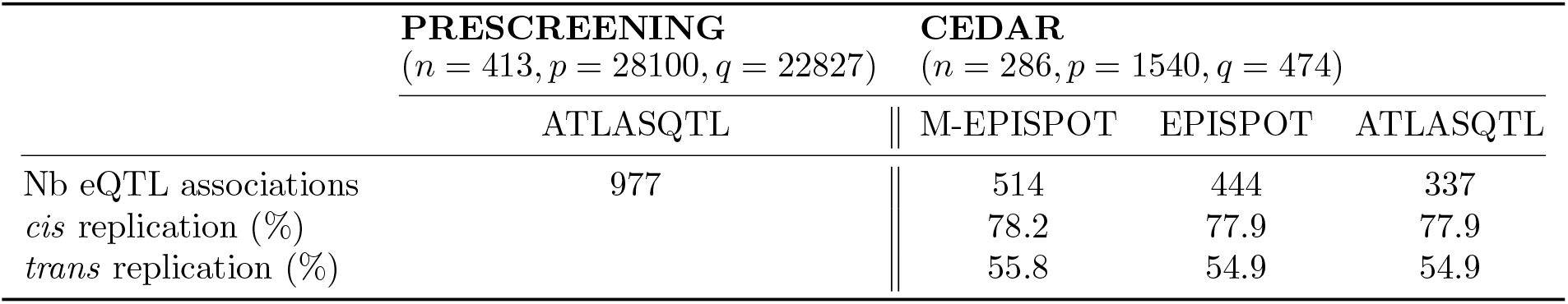
Number of hits and replication rates. Number of eQTL associations discovered by the ATLASQTL prescreening (chromosome 12) and by each of the three (M-EPISPOT, EPISPOT and ATLASQTL) analyses of the CEDAR data, along with the replication rates for the associations discovered at the prescreening stage. All analyses use an FDR threshold of 5%. The numbers of samples *n*, SNPs *p* and transcripts *q* are indicated for each dataset. Full lists of eQTL associations for the different methods are provided in Tables S1–S4.

| | **PRESCREENING**<br>($n = 413, p = 28100, q = 22827$) | **CEDAR**<br>($n = 286, p = 1540, q = 474$) | | |
| --- | --- | --- | --- | --- |
|  | ATLASQTL | \|\| M-EPISPOT | EPISPOT | ATLASQTL |
| Nb eQTL associations | 977 | \|\| 514 | 444 | 337 |
| *cis* replication (%) |  | \|\| 78.2 | 77.9 | 77.9 |
| *trans* replication (%) |  | \|\| 55.8 | 54.9 | 54.9 |

#### A focus on two susceptibility loci

We next discuss two examples of pleiotropic loci. First, not only does M-EPISPOT confirm the *LYZ* pleiotropic activity (Figure 5D-i), but it also uncovers associations of this locus with four additional genes compared to the ATLASQTL run, namely, *COPZ1, DPY30, KLHL28* and *OSTC*. The EPISPOT run (with no module partitioning) reports the exact same list as ATLASQTL, also missing the above four genes.

The second example is a pleiotropic locus uncovered by M-EPISPOT and for which only isolated effects were detected at the prescreening stage (Figure 5D-ii). This locus is located 32 Mb downstream to the *LYZ* locus and entails a hotspot of size 52 in the gene body of *GNPTAB*, namely, rs10860784 (*r*^2^ = 0.001 with the lead hotspot rs10784774 of the *LYZ* locus). The *trans* network formed by the controlled transcripts has not been previously described and nor has any *trans*-acting effect involving rs10860784 (up to proxies using *r*^2^ *>* 0.8). However rs10860784 is known to be *cis*-acting on gene *DRAM1* (located 98 Kb downstream) in multiple tissues^31^, an association which M-EPISPOT also confirms using a looser FDR of 15%. Moreover, the UK Biobank PheGWAS also reported^32^ a strong association between this SNP and height (*p* = 1.47 × 10^−14^). The module-free version EPISPOT run also finds a *trans* network for the exact same SNP, yet slightly smaller, as it involves 31 transcripts at FDR 5%; ATLASQTL finds no signal. This example suggests that the added value of epigenome-driven inference is particularly striking for the detection of weak *trans* signals. Indeed, a comparison of the estimated QTL effects attributable to rs10860784 with those attributable to *LYZ* pleiotropic locus (Figure 5D-i) shows that the former are significantly smaller in magnitude compared to the latter (*t*-test *p*-value < 2×10^−16^).

#### The selected epigenetic annotations reveal possible genetic mechanisms of action

The above figures suggest that the M-EPISPOT and EPISPOT runs allow for more powerful QTL mapping compared to ATLASQTL. This probably results from their ability to leverage the epigenetic marks, as we next discuss.

For each module, M-EPISPOT identifies a subset of epigenetic annotations with a potential to induce or inhibit the QTL associations (depending on the sign of the posterior mean of each annotation effect); these annotations are highlighted in Figure 5C. For instance, DNase-I hypersensitivity sites (DHS) in fibroblasts and epithelial cells of different tissues tend to promote the QTL effects. Interestingly, DHS in CD14^+^ monocytes are found to be enhancers of eQTL effects in both the M-EPISPOT and EPISPOT runs, with epi-PPI *>* 0.99. The two runs also estimate a negative effect of the distance to transcription start sites (TSSs, epi-PPI *>* 0.99), in line with the frequently reported decay in abundance of eQTL signals with the distance to TSS^33^. These last two observations are helpful to interpret the uncovered QTL signals, as we next discuss.

#### CD14^+^ cell DHS: hints to a monocyte-specific pleiotropic activity in LYZ

We first focus on the *LYZ* pleiotropic region. Previous studies have highlighted distinct lead hotspots around *LYZ*^34^, yet none provided a functional characterisation that would allow *a* clear prioritisation of one variant over another. The lead hotspots revealed by the M-EPISPOT and EPISPOT runs are intergenic variants, rs10784774 (size 154) and rs2168029 (size 109, *r*^2^ = 0.89 with rs10784774; see Figure 5D-i). They differ from the lead hotspot flagged by the ATLASQTL run, namely, rs1384 (size 149, *r*^2^ = 0.99 with rs10784774). We next examine the possible biology behind these candidates, starting with the ATLASQTL top hotspot.

The fact that rs1384 is located within the 3^*′*^ UTR of *LYZ* may suggest a *trans* action mediated by *LYZ*. This hypothesis is plausible given that the locus associates with *LYZ* in all M-EPISPOT, EPISPOT and ATLASQTL runs and that GTEx also reported this *cis* association in whole blood and different tissues. Conversely, regressing out the effect of *LYZ* on the expression matrix does not explain away the hotspot effects, which argues somewhat against a mediation by *LYZ* (the size of the top hotspot in the *LYZ* locus is only marginally reduced: 134 versus 154 in the original M-EPISPOT analysis, Supplemental Material and Methods).

The monocyte-specific DHS annotation selected by M-EPISPOT for the *pleiotropic module* suggests another scenario. Namely, the pleiotropic activity of the locus may be triggered by cell-type specific enhancers in open chromatin regions, which are known to be key players in activating the transcription in *trans*^35^. This hypothesis of monocyte-specific pleiotropy would also explain why no hotspot was reported so far in cell types and tissues other than monocytes^26,36^. To investigate this further, we performed an additional enrichment analysis using the multiple tissue/cell-type histone modification marks of the ENCODE catalog: we found that the two sets of genes associated with the M-EPISPOT’s lead hotspots rs10784774 and rs2168029 respectively are enriched in H3K27ac enhancers, again in CD14^+^ monocytes only, which further supports cell-type specific activation. One notable gene in this group is the transcription factor CREB1, which has previously been suggested as a putative mediator of the *LYZ* pleiotropic network^26^. Notably, regressing out the effect of *CREB1* on the expression matrix substantially reduces the pleiotropy of the locus (the size of the top hotspot in the *LYZ* locus is 36, versus 154 in the original M-EPISPOT analysis). Moreover, the connectivity of the transcript conditional independence network is also markedly lower (Supplemental Material and Methods).

We further explored whether the two sets of genes associated with either rs10784774 or rs2168029 were enriched in transcription factor binding sites (TFBS) using the ENCODE data in K562 cells. We found a profound enrichment of a number of TBFS, including ATF3, CREB1, c-Myc (Table S5). The networks of transcription factors for rs10784774 and rs2168029 are similar, indicating conserved regulatory networks.

Interestingly, ATF transcription factors are CREB-binding proteins, in line with the *CREB1* -mediation hypothesis, but the strong enrichment for many other transcription factors suggests that the same loci can be targeted by different processes and the co-occupancy of these loci in primary monocytes may resolve this further, although is important to note that, unlike the very significant association between *LYZ* and *CREB1*, there is no association between *LYZ* and *ATF3* expression, thus we can discount this gene playing a role in this genomic circuit. The c-Myc transcription factor is involved in cell division and has broad transcriptional consequences^37^, which is sensible given the large pleiotropy observed at the *LYZ* locus, for rs10784774 and rs2168029. Consistent with this, the UK Biobank data further reveal strong associations of these two SNPs with monocyte counts and other myeloid cell counts^32^.

Although by no means conclusive, these observations corroborate the context specificity of the *trans* effects controlled by the *LYZ* locus, and indeed may be more representative of other unresolved *trans* loci across the genome that, whilst of potential high biological importance, lack the pleiotropic effect of the *LYZ* locus. They also suggest that the epigenome-driven EPISPOT runs found promising candidate hotspots, whose presumed mechanisms of action on the massive *LYZ* gene network would merit experimental follow up.

#### Distance to TSSs: examples of cis and hotspot signals shared across cell types

Another interesting result concerns the negative effect of the annotation coding the distance to TSSs, this time for transcripts belonging to the *scattered module*. As active transcripts in this module are mostly involved in *cis* associations, the module-specificity of this annotation aligns with the previous observation that the distance to TSSs associates with an enrichment of *cis* eQTLs^33,38^. Moreover, an empirical assessment of this enrichment in our dataset shows that the SNPs selected with M-EPISPOT are on average significantly closer to TSSs compared to SNP subsets of the same size randomly drawn within the analysed loci (*p* = 0.017). Such an enrichment is unsurprising and actually also present in the EPISPOT and ATLASQTL results, but the importance of the distance to TSS is nevertheless made explicit by the selection of the TSS variable by both EPISPOT and M-EPISPOT.

For instance, three candidate hotspots, rs10876864, rs11171739 (*r*^2^ = 0.94 with rs10876864) and rs705699 (*r*^2^ = 0.86 with rs10876864), located 13 Mb upstream of the *LYZ* locus, are representative of this enrichment as they are within a TFBS, a 5*′* UTR and an exon, respectively (Figure 5D-iii). Our ATLASQTL prescreening and EPISPOT analyses find that they control a small network of size 11 involving transcripts mapping to the *cis* gene *RPS26* and other distal genes, including *IP6K2* on chromosome 3.

This locus has been linked with several autoimmune diseases^39–42^ including type 1 diabetes, where evidence exists that *RPS26* transcription does not mediate the disease association^43^. Interestingly, previous studies have reported the *RPS26 cis* effect as an isolated association in monocytes. The *trans* activity, in particular on *IP6K2*, was unknown in monocytes, but is known in B and T cells^26,44^. This suggests that it has so far gone unnoticed in monocytes using standard univariate mapping approaches, but our fully joint, annotation-driven method has enabled its detection. Moreover, unlike the monocyte-specific *LYZ* pleiotropic locus discussed above, this locus is an example of *trans*-hotspot eQTL present in several cell types. The genomic location also aligns with the observation that eQTLs common to multiple cell types or tissues tend to be closer to TSSs compared to eQTLs only detectable in a single cell type or tissue^45^.

## Discussion

Large panels of epigenetic marks are nowadays collected along with genetic data, and employed as part of different modelling approaches, whether for single-trait association studies or fine mapping^8–12^. However their use to enhance molecular QTL mapping remains mostly heuristic. Thanks to its hypothesis-free mark selection routine which is fully integrated within a joint QTL mapping framework, EPISPOT can tell apart the relevant epigenetic marks from thousands of candidates, while also directly refining estimation in large molecular QTL studies.

Specifically, EPISPOT brings important modelling and algorithmic contributions. First, it implements a flexible hierarchical model which enables parametrising both *cis and trans* actions on thousands of molecular traits, whereas existing epigenome-based approaches are limited to GWAS or *cis* QTL mapping for one or a handful of traits^8–12^. Second, it is both fully joint and scalable, accounting for all epigenetic marks, genetic variants and molecular levels, and their shared signals, in a single modelling framework. Third, it combines this information to perform an *automated selection of the epigenetic marks relevant to the QTL effects of the problem at hand*, thereby providing direct insight into the functional basis of the signals. Fourth, its crafted annealed variational algorithm ensures a robust exploration of complex parameters spaces, such as induced by candidate SNPs in high LD, corresponding to scenarios for which the use of epigenetic information is particularly beneficial. Finally, EPISPOT allows for module-specific learning of the epigenetic action.

We showed in a series of simulation experiments emulating epigenome-driven QTL problems that EPISPOT effectively scales to large datasets, while retaining the accuracy necessary for a powerful QTL mapping. We demonstrated that our method was not only able to pinpoint the correct marks with high posterior probability, but that it could also leverage these marks to improve the detection of weak QTL signals. In particular, we saw that the spike-and-slab representation of the epigenome contribution ensures that the irrelevant epigenetic marks are effectively discarded as “noise”, so panels with hundreds of candidate marks can be considered without the risk of worsening inferences. This allows skipping the delicate process of pre-filtering marks, whose practical grounds are often blurry and disconnected from the QTL dataset under consideration.

Our work attaches special importance to acknowledging the complexity of the learning task (selection of hotspots, pairwise QTL associations between variants and molecular traits, selection of epigenetic marks relevant to these QTL associations) and possible biological scenarios (pattern of regulation, importance of the epigenome in this regulation, dependence structures among variants, marks and molecular traits, and between them). Our simulations examined under what conditions inference is well powered to leverage the epigenetic information, and evaluated the sensitivity to different input choices, in particular when gene modules are provided. Importantly, our method is not meant to be used as a black box to fish genetic variants involved in *trans* regulation and their epigenetic roots, but rather is predicated on a careful analysis design which takes into account the dataset, the biological question of interest and the expected statistical power. Further assessments for specific problem settings (sparsity levels, association patterns and epigenetic control) can be made using the code provided online^25,46^.

Finally, we showed how our simulation studies prefigured the efficiency of EPISPOT in a large monocyte eQTL study (high replication in an independent sample, previously unreported pleiotropic loci, refined list of candidate lead hotspots). We further illustrated how the EPISPOT posterior output can be used to both select interpretable annotations underlying the QTL activity and reduce the range of hypotheses about the functional mechanisms involved, particularly for hotspots. We also showed how the localised nature of QTL activity could be accounted for when inferring annotations in a module-specific fashion using M-EPISPOT (the monocyte-specific enhancer activity affecting the *pleiotropic module*, the enrichment of QTL hits closer to TSSs affecting the *scattered module*). Altogether, this thorough case study demonstrates that QTL analyses may largely benefit from the use of rich complementary data sources annotating the primary genotyping data, provided principled joint approaches are used to capture shared association patterns.

EPISPOT offers perspectives for robust and interpretable molecular QTL mapping, towards a better understanding of the functional basis of genetic regulation. Thanks to its efficient annealed VBEM algorithm with adaptive and parallel schemes, it enables information-sharing across epigenetic marks, genetic variants and molecular traits governed by complex regulatory mechanisms, at scale. In particular, its use of selection indicators in a spike-and-slab framework allows for a systematic identification of sparse sets of epigenetic annotations which are directly relevant for the QTL regulation of the problem at hand.

We envision holistic approaches such as EPISPOT to be increasingly adopted in an age where large molecular datasets and annotation information become widely available. EPISPOT is applicable to any type of molecular QTL problem, involving genomic, proteomic, lipidomic or metabolomic levels, but also to genome-wide association with several clinical endpoints. In particular, exploiting the epigenome to build finer maps of hotspots across the genome holds great promises, as these master regulators are likely to be triggered by tissue- and cell-type-specific epigenetic functions.

## Acknowledgements

We thank Verena Zuber for her help in setting up the epigenetic annotation panel and Colin Starr for managing computational resources.

This research was funded by the UK Medical Research Council programme MRC MC UU 00002/10 (HR, SR), MC UU 00002/4 (EV, CW) and MR M0 13138/1, MR S0 2638X/1 (LB); the Engineering and Physical Sciences Research Council EP/R018561/1 (SR); the BHF-Turing Cardiovascular Data Science Awards 2017 & the Alan Turing Institute under the Engineering and Physical Sciences Research Council grant EP/N510129/1 (LB); the Alan Turing Institute Fellowship number TU/B/000092 (SR), and the Wellcome Trust WT107881 (EV, CW). This work was also supported by the NIHR Cambridge BRC. The views expressed are those of the author(s) and not necessarily those of the NHS, the NIHR or the Department of Health and Social Care. BPF and IN are funded by a Wellcome Intermediate Clinical Fellowship to BPF (no. 201488/Z/16/Z).

## Web resources

EPISPOT is implemented as an R package with C++ subroutines and is publicly available under the GNU General Public License version 3 (GPL3): https://github.com/hruffieux/epispot.

The following resources were also employed for the data processing, method comparison and eQTL analysis.

- ATLASQTL (version 0.1.4): https://github.com/hruffieux/atlasqtl
- ECHOSEQ (version 0.3.0): https://github.com/hruffieux/echoseq
- EnrichR (version 2.1): https://amp.pharm.mssm.edu/Enrichr
- Ensembl: http://grch37.ensembl.org/index.html
- GTEx: https://gtexportal.org/home
- GWAS Catalog: https://www.ebi.ac.uk/gwas
- MATRIXEQTL (version 2.3): http://www.bios.unc.edu/research/genomic_software/Matrix_eQTL
- PhenoScanner: http://www.phenoscanner.medschl.cam.ac.uk
- PLINK (version v1.90b5.3): http://zzz.bwh.harvard.edu/plink
- R (version 3.6.1): https://www.r-project.org

## Data and code availability

Fairfax et al. (2012, 2014)^26,29^ provide gene expression in CD14^+^ monocytes and genotyping data from individuals with European ancestry. The raw expression data were generated with Illumina HumanHT-12 v4 arrays and downloaded from ArrayExpress^47^ (accession E-MTAB-2232), while the raw genotyping data were generated by Illumina HumanOmniExpress-arrays and have been deposited at the European Genome-Phenome Archive (accessions: EGAD00010000144 and EGAD00010000520). The expression data are freely available, but the genotyping data require a data access agreement, as detailed in Fairfax et al. (2012, 2014)^26,29^ and https://www.well.ox.ac.uk/research/research-groups/julian-knight-group/research-projects/data-access.

The CEDAR dataset^27^ consists of gene expression data from CD14^+^ monocytes and genotyping data from individuals with European ancestry. The raw expression data were generated with Illumina HumanHT-12 v4 arrays and downloaded from ArrayExpress^47^ (accession: E-MTAB-6667), while the raw genotyping data were generated by Illumina HumanOmniExpress-12 v1 A arrays and downloaded from ArrayExpress (accession: E-MTAB-6666). Both the expression and genotyping data are freely available.

Both studies were approved by the local human research ethic committees, namely, the Oxfordshire Research Ethics Committee (COREC reference 06/Q1605/55)^29^ and the University of Liège Academic Hospital Ethics Committee^27^. Participants provided informed written consent, and all procedures were conducted in accordance with the Declaration of Helsinki.

All statistical analyses were performed using the R environment (version 3.6.1)^48^ and the synthetic datasets were generated using the freely available R package ECHOSEQ (version 0.3.0)^46^. The R package EPISPOT implements the method^25^.

## Supplemental data

- Supplemental Material and Methods.
- Table S1: Annotated 5% FDR eQTL associations using the ATLASQTL prescreening.
- Table S2: Annotated 5% FDR eQTL associations using M-EPISPOT (CEDAR) and posterior summary for epigenetic marks.
- Table S3: Annotated 5% FDR eQTL associations using EPISPOT (CEDAR) and posterior summary for epigenetic marks.
- Table S4: Annotated 5% FDR eQTL associations using ATLASQTL (CEDAR).
- Table S5: Transcription factor enrichment analysis results for the sets of genes controlled by the top hotspots.

