## Supplemental Material and Methods for "EPISPOT: an epigenome-driven approach for detecting and interpreting hotspots in molecular QTL studies"

### S.1 Hyperparameter specification for top-level priors

We describe the hyperparameter settings for the prior distribution of the response-specific parameter $\zeta_t \overset{\text{iid}}{\sim} \mathcal{N}(n_0, t_0^2)$. We let this parameter control the sparsity level, i.e., the number of predictors associated with each response, and use parameter $\theta_s \overset{\text{ind}}{\sim} \mathcal{N}(0, s_{0s}^2)$ as a predictor-specific modulator of this level.

We will rely on the following results: for $X \sim \mathcal{N}\left(\mu, \sigma^2\right)$,

$$\mathrm{E}\left\{\Phi\left(X\right)\right\} = \Phi\left(\frac{\mu}{\sqrt{1+\sigma^2}}\right), \tag{S.1}$$

$$\mathrm{E}\left\{\Phi\left(X\right)^2\right\} = \Phi\left(\frac{\mu}{\sqrt{1+\sigma^2}}\right) - 2\mathrm{T}\left(\frac{\mu}{\sqrt{1+\sigma^2}}, \frac{1}{\sqrt{1+2\sigma^2}}\right), \tag{S.2}$$

where

$$\mathrm{T}\left(h, a\right) = \varphi(h)\int_0^a \frac{\varphi(hx)}{1+x^2}\mathrm{d}x, \qquad a, h \in \mathbb{R},$$

is Owen's T function[1], with $\varphi(\cdot)$ the standard normal density function.

Equality (S.1) can be obtained as follows. Let $Z_1 \sim \mathcal{N}\left(-\sigma^{-1}\mu, \sigma^{-2}\right)$ and $Z_2 \sim \mathcal{N}\left(0,1\right)$ be independent and observe that

$$\mathrm{pr}\left(Z_1 \leq Z_2 \mid Z_2 = z\right) = \mathrm{pr}\left(Z_1 \leq z\right) = \Phi(\sigma z + \mu), \quad z \in \mathbb{R},$$

so that

$$\mathrm{pr}\left(Z_1 \leq Z_2\right) = \int \Phi(\sigma z + \mu)\varphi(z)\mathrm{d}z,$$

which corresponds to the left hand-side of (S.1). But since $Z_1 - Z_2 \sim \mathcal{N}\left(-\sigma^{-1}\mu, \sigma^{-2}+1\right)$, we also have

$$\mathrm{pr}\left(Z_1 \leq Z_2\right) = \mathrm{pr}\left(Z_1 - Z_2 \leq 0\right) = \Phi\left(\frac{\mu}{\sqrt{1+\sigma^2}}\right),$$

which gives the result. Equality (S.2) can be obtained similarly.

Coming back to the hyperparameter setting, we make the simplifying assumption that there is no predictor-specific modulation ($\theta_s = 0$) so that, given $\zeta_t$, the prior probability of association between predictor $X_s$ and response $y_t$ is

$$\mathrm{E}\left(\gamma_{st} \mid \theta_s = 0, \zeta_t\right) = \Phi(\zeta_t).$$

We then set $n_0$ and $t_0^2$ by specifying a prior expectation and a prior variance for the number of predictors associated with each response, $p_{\gamma,t} = \sum_{s=1}^{p}\gamma_{st}$, $t = 1, \ldots, q$,

$$\begin{aligned}
\mathrm{E}\left(p_{\gamma,t} \mid \theta = 0\right) &= \mathrm{E}\left\{\mathrm{E}\left(p_{\gamma,t} \mid \theta = 0, \zeta_t\right)\right\} = p\,\mathrm{E}\left\{\Phi\left(\zeta_t\right)\right\}, \\
\mathrm{Var}\left(p_{\gamma,t} \mid \theta = 0\right) &= \mathrm{Var}\left\{\mathrm{E}\left(p_{\gamma,t} \mid \theta = 0, \zeta_t\right)\right\} + \mathrm{E}\left\{\mathrm{Var}\left(p_{\gamma,t} \mid \theta = 0, \zeta_t\right)\right\} \\
&= p(p-1)\mathrm{E}\left\{\Phi\left(\zeta_t\right)^2\right\} + p\mathrm{E}\left\{\Phi\left(\zeta_t\right)\right\}\left[1 - p\mathrm{E}\left\{\Phi\left(\zeta_t\right)\right\}\right],
\end{aligned}$$

in which we use (S.1) and (S.2) with $\mu = n_0$ and $\sigma^2 = t_0^2$. We then solve this system numerically to obtain $n_0$ and $t_0^2$.

### S.2 Derivation of the variational expectation-maximisation algorithm

**S.2.1 Variational distributions.** We provide here the detailed derivation of the variational expectation-maximisation (VBEM) algorithm. We describe the algorithm in its general module-based form (M-EPISPOT); omitting the index $m$ and taking $M = 1$ gives the base version with no module partitioning (EPISPOT). Let $\boldsymbol{y} = (\boldsymbol{y}_1, \ldots, \boldsymbol{y}_q)$ be an $n \times q$ matrix of $q$ centred responses, $\boldsymbol{X} = (\boldsymbol{X}_1, \ldots, \boldsymbol{X}_p)$ be an $n \times p$ matrix of $p$ centred predictors, for $n$ samples, and $\boldsymbol{V} = (V_1, \ldots V_r)$ is a $p \times r$ matrix of $r$ centred predictor-level covariates. We rewrite model (1)–(3) using the classical data-augmentation representation of the probit-link level, i.e., by introducing the auxiliary variable $z_{st}$ as follows:

$$\begin{array}{rcllr} \boldsymbol{y}_t \mid \boldsymbol{\beta}_t, \tau_t & \sim & \mathcal{N}_n\left(\boldsymbol{X}\boldsymbol{\beta}_t, \tau_t^{-1}\boldsymbol{I}_n\right), & \tau_t \sim \text{Gamma}(\eta_t, \kappa_t), & t = 1, \ldots, q, \\ \beta_{st} \mid \gamma_{st}, \sigma^2, \tau_t & \sim & \gamma_{st}\mathcal{N}\left(0, \sigma^2\tau_t^{-1}\right) + (1-\gamma_{st})\,\delta_0, & \sigma^{-2} \sim \text{Gamma}(\lambda, \nu), & s = 1, \ldots, p, \\ \gamma_{st} & = & \mathbb{1}\{z_{st} > 0\}, & z_{st} \mid \theta_{m,s}, \zeta_t, \boldsymbol{\xi}_m \sim \mathcal{N}\left(\theta_{m,s} + \zeta_t + \boldsymbol{V}_s^T\boldsymbol{\xi}_m, 1\right), & \\ \xi_{m,l} \mid \rho_{m,l} & \sim & \rho_{m,l}\mathcal{N}\left(0, s_m^2\right) + (1-\rho_{m,l})\,\delta_0, & \theta_{m,s} \sim \mathcal{N}(0, s_{0m,s}^2), & \zeta_t \sim \mathcal{N}(n_0, t_0^2), \\ \rho_{m,l} & \sim & \text{Bernoulli}\left(\omega_{m,l}\right), & & l = 1, \ldots, r, \end{array}$$

where $m \in \mathcal{M}$ is a module of response variables, with $\mathcal{M}$ a partition of $\{1, \ldots, q\}$, $m \ni t$.

Let $\boldsymbol{v} = \left(\boldsymbol{\beta}, \boldsymbol{\gamma}, \boldsymbol{z}, \boldsymbol{\theta}, \boldsymbol{\zeta}, \boldsymbol{\xi}, \boldsymbol{\rho}, \boldsymbol{\tau}, \sigma^{-2}\right)$ be the parameter vector, $\boldsymbol{\eta}_m = \left(\boldsymbol{s}_{0m}^2, s_m^2, \boldsymbol{\omega}_m\right)$ be the hyper-parameter vector for the second-stage model for module $m$, and $\boldsymbol{\eta} = (\boldsymbol{\eta}_1, \ldots, \boldsymbol{\eta}_M)$. We have

$$\begin{aligned} p(\boldsymbol{y}, \boldsymbol{v} \mid \boldsymbol{\eta}) \;=\; & \left\{\prod_{t=1}^q p\left(\boldsymbol{y}_t \mid \boldsymbol{\beta}_t, \tau_t\right)\right\}\left\{\prod_{t=1}^q\prod_{s=1}^p p\left(\beta_{st} \mid \gamma_{st}, \sigma^{-2}, \tau_t\right)\right\}\left\{\prod_{t=1}^q p\left(\tau_t\right)\right\} p(\sigma^{-2}) \\ & \times \left\{\prod_{t=1}^q\prod_{s=1}^p p\left(\gamma_{st} \mid z_{st}\right) p(z_{st} \mid \theta_{m,s}, \zeta_t, \boldsymbol{\xi}_m)\right\}\left\{\prod_{m=1}^M\prod_{s=1}^p p(\theta_{m,s} \mid s_{0m,s}^2)\right\} \\ & \times \left\{\prod_{t=1}^q p(\zeta_t)\right\}\left\{\prod_{m=1}^M\prod_{l=1}^r p(\xi_{m,l} \mid \rho_{m,l}, s_m^2) p(\rho_{m,l} \mid \omega_{m,l})\right\}, \end{aligned}$$

where, in the second conditional distribution of the second line $p(z_{st} \mid \theta_{m,s}, \zeta_t, \boldsymbol{\xi}_m)$, $m = m(t)$ implicitly corresponds to the module containing response $\boldsymbol{y}_t$; we adopt such tacit notation hereafter for brevity.

We use the following mean-field form for the variational approximation,

$$\begin{aligned} q\left(\boldsymbol{v}\right) \;=\; & \left\{\prod_{t=1}^q\prod_{s=1}^p q(\beta_{st}, \gamma_{st}, z_{st})\right\}\left\{\prod_{t=1}^q q\left(\tau_t\right)\right\} q\left(\sigma^{-2}\right)\left\{\prod_{m=1}^M\prod_{s=1}^p q(\theta_{m,s})\right\}\left\{\prod_{t=1}^q q(\zeta_t)\right\} \\ & \times \left\{\prod_{m=1}^M\prod_{l=1}^r q(\xi_{m,l}, \rho_{m,l})\right\}, \end{aligned}$$

and augment the variational algorithm with annealing steps in the first iterations. Namely, we introduce a *temperature parameter* $T \geq 1$ and, writing $q_T(\boldsymbol{v})$ the *heated* variational approximation, we maximize the following annealed variational objective function:

$$\mathcal{L}_T(q_T) = \int q_T(\boldsymbol{v}) \log p(\boldsymbol{v}, \boldsymbol{y}) \mathrm{d}v - T \int q_T(\boldsymbol{v}) \log q_T(\boldsymbol{v}) \mathrm{d}\boldsymbol{v}\,.$$

We derive the form of the heated variational distribution $q_T(v_j)$ by observing that

$$\begin{aligned}
\mathcal{L}_T(q_T) &= \mathrm{E}_j\left[\mathrm{E}_{-j}\left\{\log p(\boldsymbol{v},\boldsymbol{y})\right\} - T\log q_T(v_j)\right] + \mathrm{cst} \\
&= \mathrm{E}_j\left[\log\left\{\frac{\exp\left\{\mathrm{E}_{-j}\log p(\boldsymbol{v},\boldsymbol{y})\right\}}{q_T(v_j)^T}\right\}\right] + \mathrm{cst} \\
&= T\,\mathrm{E}_j\left[\log\left\{\frac{p_{T,-j}(v_j,\boldsymbol{y})}{q_T(v_j)}\right\}\right] + \mathrm{cst}, \qquad \text{(S.3)}
\end{aligned}$$

where we introduced the distribution $p_{T,-j}(v_j,\boldsymbol{y}) \propto \exp\left\{T^{-1}\mathrm{E}_{-j}\log p(\boldsymbol{v},\boldsymbol{y})\right\}$, and where $\mathrm{E}_j(\cdot)$ denotes the expectation with respect to the distribution $q_T(v_j)$, $\mathrm{E}_{-j}(\cdot)$, the expectation with respect to the distributions $q_T(v_k)$, for all the variables $v_k$ $(k \neq j)$, and cst is constant with respect to $v_j$. The expectation in (S.3) corresponds to the negative Kullback–Leibler divergence between $q_T(v_j)$ and the $p_{T,-j}(v_j,\boldsymbol{y})$; $\mathcal{L}_T(q)$ is therefore maximal when $q_T(v_j) = p_{T,-j}(v_j,\boldsymbol{y})$, i.e., when

$$\log q_T(v_j) = T^{-1}\mathrm{E}_{-j}\{\log p(\boldsymbol{y},\boldsymbol{v})\} + \mathrm{cst}\,, \qquad j = 1,\ldots,J\,. \qquad \text{(S.4)}$$

For ease of reading, we hereafter drop the subscript $T$ in $q_T(\cdot)$, and write $c = T^{-1}$ and $v_j^{(r)}$ for the $r^{th}$ moment with respect to the approximate posterior distribution $q(v_j)$. We find that,

$$q(\beta_{st},\gamma_{st},z_{st}) = q(\beta_{st}\mid z_{st})q(z_{st}\mid\gamma_{st})q(\gamma_{st})\,, \qquad s = 1,\ldots,p,\ t = 1,\ldots,q$$

with

$$\begin{aligned}
\beta_{st}\mid z_{st} > 0,\boldsymbol{y} &\sim \mathcal{N}\left(\mu_{\beta,st},\sigma^2_{\beta,st}\right)\,, \quad \beta_{st}\mid z_{st}\leq 0,\boldsymbol{y} \sim \delta_0\,, \\
z_{st}\mid\gamma_{st} = \delta,\boldsymbol{y} &\sim \mathcal{TN}\left(\theta^{(1)}_{m,s} + \zeta^{(1)}_t + \boldsymbol{V}_s^T\boldsymbol{\xi}^{(1)}_m, c^{-1}; \left\{0 < (-1)^{1-\delta}z_{st}\right\}\right)\,,\ \delta = 0,1, \\
\gamma_{st}\mid\boldsymbol{y} &\sim \mathrm{Bernoulli}\left(\gamma^{(1)}_{st}\right)\,,
\end{aligned}$$

where $X \sim \mathcal{TN}\left(\mu,\sigma^2;\{a < x < b\}\right)$ denotes a truncated normal variable,

$$\sigma^{-2}_{\beta,st} = c\,\tau^{(1)}_t\left\{\|\boldsymbol{X}_s\|^2 + \left(\sigma^{-2}\right)^{(1)}\right\}\,, \qquad \mu_{\beta,st} = c\,\sigma^2_{\beta,st}\tau^{(1)}_t\boldsymbol{X}_s^T\left(\boldsymbol{y}_t - \sum_{j=1,j\neq s}^{p}\gamma^{(1)}_{jt}\mu_{\beta,jt}\boldsymbol{X}_j\right)\,,$$

and

$$\begin{aligned}
\frac{1}{\gamma^{(1)}_{st}} &= 1 + \exp\left[-c\left\{\frac{1}{2}\left(\log\sigma^{-2}\right)^{(1)} + \frac{1}{2}\left(\log\tau_t\right)^{(1)} + \frac{1}{2}\mu^2_{\beta,st}\sigma^{-2}_{\beta,st} + \log\sigma_{\beta,st}\right.\right. \\
&\qquad \left.\left. - \log\left\{1 - \Phi\left(\theta^{(1)}_{m,s} + \zeta^{(1)}_t + \boldsymbol{V}_s^T\boldsymbol{\xi}^{(1)}_m\right)\right\} + \log\Phi\left(\theta^{(1)}_{m,s} + \zeta^{(1)}_t + \boldsymbol{V}_s^T\boldsymbol{\xi}^{(1)}_m\right)\right\}\right]\,.
\end{aligned}$$

Writing $\alpha_{st} = \theta_{m,s} + \zeta_t + \boldsymbol{V}_s^T\boldsymbol{\xi}_m$, the first moment of $z_{st}$ given $\gamma_{st}$ is

$$\mathrm{E}_q\left(z_{st}\mid\gamma_{st}\right) = \alpha^{(1)}_{st} + c^{-1/2}M\left(c^{1/2}\alpha^{(1)}_{st},\gamma_{st}\right)\,,$$

where

$$M(u,\gamma) = (-1)^{1-\gamma}\frac{\varphi(u)}{\Phi(u)^\gamma\left[1 - \Phi(u)\right]^{1-\gamma}}, \qquad u\in\mathbb{R},\ \gamma = 0,1,$$

is the inverse Mills ratio and $\mathrm{E}_q(\cdot)$ is the expectation with respect to the variational distribution $q(\cdot)$. We therefore have

$$\begin{aligned}
z^{(1)}_{st} &= \gamma^{(1)}_{st}\left(\alpha^{(1)}_{st} + c^{-1/2}M(c^{1/2}\alpha^{(1)}_{st},1)\right) + (1 - \gamma^{(1)}_{st})\left(\alpha^{(1)}_{st} + c^{-1/2}M(c^{1/2}\alpha^{(1)}_{st},0)\right) \\
&= c^{-1/2}\gamma^{(1)}_{st}\left\{M(c^{1/2}\alpha^{(1)}_{st},1) - M(c^{1/2}\alpha^{(1)}_{st},0)\right\} + \alpha^{(1)}_{st} + c^{-1/2}M(c^{1/2}\alpha^{(1)}_{st},0).
\end{aligned}$$

The second moment of $z_{st}$ given $\gamma_{st}$ is

$$\begin{aligned}\mathrm{E}_q\left(z_{st}^2 \mid \gamma_{st}\right) &= c^{-1} + \left(\alpha_{st}^{(1)}\right)^2 - c^{-1/2}\alpha_{st}^{(1)} M\left(c^{1/2}\alpha_{st}^{(1)}, \gamma_{st}\right) + 2c^{-1/2}\alpha_{st}^{(1)} M\left(c^{1/2}\alpha_{st}^{(1)}, \gamma_{st}\right) \\ &= c^{-1} + \alpha_{st}^{(1)}\mathrm{E}_q\left(z_{st} \mid \gamma_{st}\right),\end{aligned}$$

which implies that

$$\begin{aligned}z_{st}^{(2)} &= c^{-1}\gamma_{st}^{(1)} + \gamma_{st}^{(1)}\alpha_{st}^{(1)}\mathrm{E}_q\left(z_{st} \mid \gamma_{st}=1\right) + c^{-1}(1-\gamma_{st}^{(1)}) + (1-\gamma_{st}^{(1)})\alpha_{st}^{(1)}\mathrm{E}_q\left(z_{st} \mid \gamma_{st}=0\right) \\ &= c^{-1} + \alpha_{st}^{(1)} z_{st}^{(1)},\end{aligned}$$

and finally its entropy is

$$H(z_{st} \mid \gamma_{st}) = \log\left[\sqrt{\frac{2\pi e}{c}}\,\Phi\left(c^{1/2}\alpha_{st}^{(1)}\right)^{\gamma_{st}}\left\{1-\Phi\left(c^{1/2}\alpha_{st}^{(1)}\right)\right\}^{1-\gamma_{st}}\right] - \frac{1}{2}c^{1/2}\alpha_{st}^{(1)} M\left(c^{1/2}\alpha_{st}^{(1)}, \gamma_{st}\right). \tag{S.5}$$

Then, we find

$$\sigma^{-2} \mid \boldsymbol{y} \sim \mathrm{Gamma}\left(\nu_\sigma, \rho_\sigma\right), \qquad \left(\sigma^{-2}\right)^{(1)} = \nu_\sigma/\rho_\sigma,$$

with

$$\nu_\sigma = c\left(\nu + \frac{1}{2}\sum_{t=1}^{q}\sum_{s=1}^{p}\gamma_{st}^{(1)}\right) - c + 1, \qquad \rho_\sigma = c\left\{\rho + \frac{1}{2}\sum_{t=1}^{q}\sum_{s=1}^{p}\gamma_{st}^{(1)}\left(\mu_{\beta,st}^2 + \sigma_{\beta,st}^2\right)\tau_t^{(1)}\right\}.$$

The residual precision parameters have

$$\tau_t \mid \boldsymbol{y} \sim \mathrm{Gamma}\left(\eta_{\tau,t}, \kappa_{\tau,t}\right), \qquad \tau_t^{(1)} = \eta_{\tau,t}/\kappa_{\tau,t}\,,$$

where

$$\begin{aligned}\eta_{\tau,t} &= c\left(\eta_t + \frac{n}{2} + \frac{1}{2}\sum_{s=1}^{p}\gamma_{st}^{(1)}\right) - c + 1\,, \\ \kappa_{\tau,t} &= c\left[\kappa_t + \frac{1}{2}\|\boldsymbol{y}_t\|^2 - \boldsymbol{y}_t^T\sum_{s=1}^{p}\mu_{\beta,st}\gamma_{st}^{(1)}\boldsymbol{X}_s + \sum_{s=1}^{p-1}\mu_{\beta,st}\gamma_{st}^{(1)}\boldsymbol{X}_s^T\sum_{j=s+1}^{p}\mu_{\beta,jt}\gamma_{jt}^{(1)}\boldsymbol{X}_j\right. \\ &\qquad \left. + \frac{1}{2}\sum_{s=1}^{p}\gamma_{st}^{(1)}\left(\sigma_{\beta,st}^2 + \mu_{\beta,st}^2\right)\left\{\|\boldsymbol{X}_s\|^2 + \left(\sigma^{-2}\right)^{(1)}\right\}\right].\end{aligned}$$

We then have

$$\theta_{m,s} \mid \boldsymbol{y} \sim \mathcal{N}\left(\mu_{\theta,m,s}, \sigma_{\theta,m,s}^2\right),$$

with

$$\sigma_{\theta,m,s}^{-2} = c\left(q_m + s_{0m,s}^{-2}\right), \qquad \mu_{\theta,m,s} = c\,\sigma_{\theta,m,s}^2\left\{\sum_{t\in m}\left(z_{st}^{(1)} - \zeta_t^{(1)}\right) - \boldsymbol{V}_s^T\boldsymbol{\xi}_m^{(1)}\right\},$$

where $q_m$ is the number of responses in module $m$, $t \in m$ means $t$ is a response index from module $m$. For $\zeta_t$, we find

$$\zeta_t \mid \boldsymbol{y} \sim \mathcal{N}\left(\mu_{\zeta,t}, \sigma_{\zeta,t}^2\right),$$

with

$$\sigma_{\zeta,t}^{-2} = c\left(p + t_0^{-2}\right), \qquad \mu_{\zeta,t} = c\,\sigma_{\zeta,t}^2\left\{\sum_{s=1}^{p}\left(z_{st}^{(1)} - \theta_{m,s}^{(1)} - \boldsymbol{V}_s^T\boldsymbol{\xi}_m^{(1)}\right) + t_0^{-2}n_0\right\}.$$

The variational distribution for the effects of the predictor-level covariates is

$$q(\xi_{m,l}, \rho_{m,l}) = q(\xi_{m,l} \mid \rho_{m,l}) q(\rho_{m,l}),$$

with

$$\xi_{m,l} \mid \rho_{m,l} = 1, \boldsymbol{y} \sim \mathcal{N}\left(\mu_{\xi,m,l}, \sigma^2_{\xi,m,l}\right), \quad \xi_{m,l} \mid \rho_{m,l} = 0, \boldsymbol{y} \sim \delta_0, \quad \rho_{m,l} \mid \boldsymbol{y} \sim \text{Bernoulli}\left(\rho^{(1)}_{m,l}\right),$$

where

$$\sigma^{-2}_{\xi,m,l} = c\left\{q_m \sum_{s=1}^{p} V^2_{sl} + s^{-2}_m\right\},$$

$$\mu_{\xi,m,l} = c\sigma^2_{\xi,m,l} \sum_{s=1}^{p} V_{sl} \left\{\sum_{t\in m} z_{st} - q_m \mu_{\theta,m,s} - \sum_{t \in m} \mu_{\zeta,t} - q_m \sum_{j=1, j\neq l}^{r} \rho^{(1)}_{m,j} \mu_{\xi,m,j} V_{sj}\right\},$$

and

$$\frac{1}{\rho^{(1)}_{m,l}} = 1 + \exp\left[-c\left\{\log \omega_{m,l} + \frac{1}{2}\mu^2_{\xi,m,l}\sigma^{-2}_{\xi,m,l} - \frac{1}{2}\log s^2_m - \log(1-\omega_{m,l}) + \log \sigma_{\xi,m,l}\right\}\right].$$

**S.2.2 Variational lower bound.** We now provide the computational details for the lower bound, $\mathcal{L}(q; \boldsymbol{\eta})$, of the marginal log-likelihood, $\log p(\boldsymbol{y} \mid \boldsymbol{\eta})$. $\mathcal{L}(q)$ is used to monitor convergence and evaluated adaptively with the convergence status for efficient saving of the computational resources. Namely, it evaluated only after the final temperature $T = 1$ is reached and not at each iteration but after a certain number of iterations which is progressively reduced as the changes in $\mathcal{L}(q)$ approaches the tolerance.

$$\begin{aligned}
\mathcal{L}(q;\boldsymbol{\eta}) &= \int q(\boldsymbol{v}) \log\left\{\frac{p(\boldsymbol{y},\boldsymbol{v})}{q(\boldsymbol{v})}\right\} \mathrm{d}\boldsymbol{v} \\
&= \sum_{t=1}^{q} \mathcal{L}_y\left(\boldsymbol{y}_t \mid \boldsymbol{\beta}_t, \boldsymbol{\gamma}_t, \tau_t\right) + \sum_{t=1}^{q}\sum_{s=1}^{p} \mathcal{L}_{\beta,\gamma}\left(\beta_{st}, \gamma_{st}, z_{st} \mid \sigma^{-2}, \tau_t, \theta_{m,s}, \zeta_t, \boldsymbol{\xi}_m\right) + \sum_{t=1}^{q} \mathcal{L}_\tau(\tau_t) \qquad \text{(S.6)} \\
&\quad + \mathcal{L}_\sigma\left(\sigma^{-2}\right) + \sum_{m=1}^{M}\sum_{s=1}^{p} \mathcal{L}_\theta\left(\theta_{m,s} \mid s^2_{0m,s}\right) + \sum_{t=1}^{q} \mathcal{L}_\zeta(\zeta_t) + \sum_{m=1}^{M}\sum_{l=1}^{r} \mathcal{L}_{\xi,\rho}\left(\xi_{m,l}, \rho_{m,l} \mid s^2_m, \omega_{m,l}\right),
\end{aligned}$$

where we recall the tacit notation $t \in m(t) = m$ and

$$\begin{aligned}
\mathcal{L}_y\left(\boldsymbol{y}_t \mid \boldsymbol{\beta}_t, \boldsymbol{\gamma}_t, \tau_t\right) &= \mathrm{E}_q\left\{\log p(\boldsymbol{y}_t \mid \boldsymbol{\beta}_t, \boldsymbol{\gamma}_t, \tau_t)\right\} \\
&= -\frac{n}{2}\log(2\pi) + \frac{n}{2}\mathrm{E}(\log \tau_t) - \tau^{(1)}_t\left\{\kappa_{\tau,t} - \frac{1}{2}\sum_{s=1}^{p} \gamma^{(1)}_{st}\left(\sigma^2_{\beta,st} + \mu^2_{\beta,st}\right)\left(\sigma^{-2}\right)^{(1)} - \kappa_t\right\},
\end{aligned}$$

$$\begin{aligned}
\mathcal{L}_{\beta,\gamma}\left(\beta_{st}, \gamma_{st}, z_{st} \mid \sigma^{-2}, \tau_t, \theta_{m,s}, \zeta_t, \boldsymbol{\xi}_m\right) &= \mathrm{E}_q \log p\left(\beta_{st} \mid \gamma_{st}, \sigma^{-2}, \tau_t\right) + \mathrm{E}_q \log p\left(\gamma_{st} \mid z_{st}\right) \\
&\quad + \mathrm{E}_q \log p\left(z_{st} \mid \theta_{m,s}, \zeta_t, \boldsymbol{\xi}_m\right) - \mathrm{E}_q \log q\left(\beta_{st}, \gamma_{st}, z_{st}\right) \\
&= \frac{1}{2}\gamma^{(1)}_{st}\left\{\mathrm{E}_q\left(\log \sigma^{-2}\right) + \mathrm{E}_q(\log \tau_t) - \left(\mu^2_{\beta,st} + \sigma^2_{\beta,st}\right)\left(\sigma^{-2}\right)^{(1)} \tau^{(1)}_t\right\} \\
&\quad + \frac{1}{2}\gamma^{(1)}_{st}\left(\log \sigma^2_{\beta,st} + 1\right) + \gamma^{(1)}_{st} \log \Phi\left(\theta^{(1)}_{m,s} + \zeta^{(1)}_t + \boldsymbol{V}_s^T \boldsymbol{\xi}^{(1)}_m\right) \\
&\quad + \left(1 - \gamma^{(1)}_{st}\right) \log\left\{1 - \Phi\left(\theta^{(1)}_{m,s} + \zeta^{(1)}_t + \boldsymbol{V}_s^T \boldsymbol{\xi}^{(1)}_m\right)\right\} \\
&\quad - \frac{1}{2}\sigma^2_{\theta,m,s} - \frac{1}{2}\sigma^2_{\zeta,t} - \frac{1}{2}\sum_{l=1}^{r} V^2_{sl}\left\{\sigma^2_{\xi,l} + \mu^2_{\xi,l}(1 - \rho^{(1)}_l)\right\}\rho^{(1)}_l \\
&\quad - \gamma^{(1)}_{st} \log \gamma^{(1)}_{st} - \left(1 - \gamma^{(1)}_{st}\right) \log\left(1 - \gamma^{(1)}_{st}\right),
\end{aligned}$$

$$\begin{aligned}\mathcal{L}_{\tau}\left(\tau_t\right) &= \mathrm{E}_q\left\{\log p(\tau_t)\right\} - \mathrm{E}_q\left\{\log q(\tau_t)\right\} \\ &= \left(\eta_t - \eta_{\tau,t}\right)\left(\log \tau_t\right)^{(1)} - \left(\kappa_t - \kappa_{\tau,t}\right)\tau_t^{(1)} + \eta_t \log \kappa_t - \eta_{\tau,t}\log \kappa_{\tau,t} - \log \Gamma(\eta_t) + \log \Gamma(\eta_{\tau,t}),\end{aligned}$$

and

$$\begin{aligned}\mathcal{L}_{\sigma}\left(\sigma^{-2}\right) &= \mathrm{E}_q \log p\left(\sigma^{-2}\right) - \mathrm{E}_q \log q\left(\sigma^{-2}\right) \\ &= \left(\nu - \nu_{\sigma}\right)\left(\log \sigma^{-2}\right)^{(1)} - \left(\rho - \rho_{\sigma}\right)\left(\sigma^{-2}\right)^{(1)} + \nu \log \rho - \nu_{\sigma}\log \rho_{\sigma} - \log \Gamma\left(\nu\right) + \log \Gamma\left(\nu_{\sigma}\right).\end{aligned}$$

We then find

$$\begin{aligned}\mathcal{L}_{\theta}(\theta_{m,s} \mid s^2_{0m,s}) &= \mathrm{E}_q\left\{\log p(\theta_{m,s} \mid s^2_{0m,s})\right\} - \mathrm{E}_q\left\{\log q(\theta_{m,s})\right\} \\ &= \frac{1}{2}\left\{-\log s^2_{0m,s} + \log \sigma^2_{\theta,m,s} - s^{-2}_{0m,s}\left(\mu^2_{\theta,m,s} + \sigma^2_{\theta,m,s}\right) + 1\right\},\end{aligned}$$

$$\begin{aligned}\mathcal{L}_{\zeta}\left(\zeta_t\right) &= \mathrm{E}_q\left\{\log p(\zeta_t)\right\} - \mathrm{E}_q\left\{\log q(\zeta_t)\right\} \\ &= \frac{1}{2}\left\{-\log t_0^2 + \log\left(\sigma^2_{\zeta,t}\right) - t_0^{-2}\left(\mu_{\zeta,t} - n_0\right)^2 - t_0^{-2}\sigma^2_{\zeta,t} + 1\right\},\end{aligned}$$

and

$$\begin{aligned}\mathcal{L}_{\xi,\rho}\left(\xi_{m,l}, \rho_{m,l} \mid s^2_m, \omega_{m,l}\right) &= -\frac{1}{2}\rho^{(1)}_{m,l}\log s^2_m + \frac{1}{2}\rho^{(1)}_{m,l}\left(\log \sigma^2_{\xi,m,l} + 1\right) - \rho^{(1)}_{m,l}\log \rho^{(1)}_{m,l} - (1 - \rho^{(1)}_{m,l})\log\left(1 - \rho^{(1)}_{m,l}\right) \\ &\quad - \frac{1}{2s^2_m}\rho^{(1)}_{m,l}\left(\sigma^2_{\xi,m,l} + \mu^2_{\xi,m,l}\right) + \rho^{(1)}_{m,l}\log \omega_{m,l} + \left(1 - \rho^{(1)}_{m,l}\right)\log\left(1 - \omega_{m,l}\right).\end{aligned}$$

**S.2.3 EM hyperparameter updates.** The M-step updates of the VBEM algorithm are obtained by taking the first derivative of (S.6) with respect to the hyperparameters. We have

$$\frac{\partial}{\partial s^2_{0m,s}}\mathcal{L}(q;\boldsymbol{\eta}) = \frac{\partial}{\partial s^2_{0m,s}}\mathcal{L}_{\theta}(\theta_{m,s} \mid s^2_{0m,s}) = -\frac{1}{2}s^{-2}_{0m,s} + \frac{1}{2}s^{-4}_{0m,s}\left(\mu^2_{\theta,m,s} + \sigma^2_{\theta,m,s}\right),$$

so

$$s^2_{0m} = \mu^2_{\theta,m,s} + \sigma^2_{\theta,m,s},$$

$$\frac{\partial}{\partial s^2_m}\mathcal{L}(q;\boldsymbol{\eta}) = \frac{\partial}{\partial s^2_m}\sum_{l=1}^{r}\mathcal{L}_{\xi,\rho}(\xi_{m,l}, \rho_{m,l} \mid s^2_m, \omega_{m,l}) = -\frac{1}{2}s_m^{-2}\sum_{l=1}^{r}\rho^{(1)}_{m,l} + \frac{1}{2}s_m^{-4}\sum_{l=1}^{r}\rho^{(1)}_{m,l}\left(\mu^2_{\xi,m,l} + \sigma^2_{\xi,m,l}\right),$$

so

$$s^2_m = \frac{\sum_{l=1}^{r}\rho^{(1)}_{m,l}\left(\mu^2_{\xi,m,l} + \sigma^2_{\xi,m,l}\right)}{\sum_{l=1}^{r}\rho^{(1)}_{m,l}},$$

and finally,

$$\frac{\partial}{\partial \omega_{m,l}}\mathcal{L}(q;\boldsymbol{\eta}) = \frac{\partial}{\partial \omega_{m,l}}\mathcal{L}_{\xi,\rho}(\xi_{m,l}, \rho_{m,l} \mid s^2_m, \omega_{m,l}) = \frac{\rho^{(1)}_{m,l}}{\omega_{m,l}} - \frac{1 - \rho^{(1)}_{m,l}}{1 - \omega_{m,l}},$$

so

$$\omega_{m,l} = \rho^{(1)}_{m,l}.$$

**S.2.4 Algorithm.** We provide a sketch of the EPISPOT VBEM algorithm. For brevity, the coupling with simulated annealing is not described, but it applies to both the E-step and the final variational run.

**Algorithm 1:** EPISPOT VBEM algorithm

**Define:** Parameters $\boldsymbol{v} = (v_1, \ldots, v_J)$, hyperparameters $\boldsymbol{\eta} = (\boldsymbol{\eta}_1, \ldots, \boldsymbol{\eta}_M)$, $\boldsymbol{\eta}_m = (\boldsymbol{s}_{0m}^2, s_m^2, \boldsymbol{\omega}_m)$.

**1. VBEM runs for hyperparameter estimation**

**for** *m = 1, ..., M (parallel loop)* **do**

  **Input:** Responses for module $m$, predictors and predictor-level covariates: $\boldsymbol{y}_m$, $\boldsymbol{X}$, $\boldsymbol{V}$

  **Output:** Empirical Bayes hyperparameter estimate: $\hat{\boldsymbol{\eta}}_m$

  **initialise**: $\boldsymbol{\eta}_m^{(0)}$, $t \leftarrow 0$

  **repeat**

    $t \leftarrow t + 1$

    <u>**E-step**</u>:

    **Input:** Current hyperparameter value: $\boldsymbol{\eta}_m^{(t-1)}$

    **Output:** Top-level model variational parameters: $\boldsymbol{\mu}_\theta$, $\boldsymbol{\sigma}_\theta^2$, $\boldsymbol{\mu}_\xi$, $\boldsymbol{\sigma}_\xi^2$, $\boldsymbol{\rho}^{(1)}$ (dropping label $m$)

    **repeat**

      **for** $j = \text{shuffle}(1, \ldots, J_m)$ **do**

        $q_m\left(v_j; \boldsymbol{\eta}_m^{(t-1)}\right) \propto \exp\left\{\mathrm{E}_{-j} \log p\left(\boldsymbol{v}, \boldsymbol{y}_m \mid \boldsymbol{\eta}_m^{(t-1)}\right)\right\}$,

      **end**

    **until** convergence of all variational parameters (with adaptive tolerance);

    <u>**M-step**</u>:

    **Input:** Current variational parameter values: $\boldsymbol{\mu}_\theta$, $\boldsymbol{\sigma}_\theta^2$, $\boldsymbol{\mu}_\xi$, $\boldsymbol{\sigma}_\xi^2$, $\boldsymbol{\rho}^{(1)}$

    **Output:** Updated hyperparameter value: $\boldsymbol{\eta}_m^{(t)}$

    $s_{0m,s}^2 \leftarrow \mu_{\theta,s}^2 + \sigma_{\theta,s}^2, \quad s = 1, \ldots, p,$

    $s_m^2 \leftarrow \dfrac{\sum_{l=1}^r \rho_l^{(1)} \left(\mu_{\xi,l}^2 + \sigma_{\xi,l}^2\right)}{\sum_{l=1}^r \rho_l^{(1)}}$

    $\omega_{m,l} \leftarrow \rho_l^{(1)}, \quad l = 1, \ldots, r,$

    $\boldsymbol{\eta}_m^{(t)} \leftarrow (\boldsymbol{s}_{0m}^2, s_m^2, \boldsymbol{\omega}_m)$

  **until** convergence of $\boldsymbol{\eta}_m^{(t)}$;

  $\hat{\boldsymbol{\eta}}_m \leftarrow \boldsymbol{\eta}_m^{(t)}$

**end**

**2. Final variational run**

**Input:** All responses, predictors and predictor-level covariates: $\boldsymbol{y}$, $\boldsymbol{X}$, $\boldsymbol{V}$, empirical Bayes hyperparameter: $\hat{\boldsymbol{\eta}}$

**Output:** Variational parameters

**repeat**

  **for** $j = \text{shuffle}(1, \ldots, J)$ **do**

    $q\left(v_j; \hat{\boldsymbol{\eta}}\right) \propto \exp\left\{\mathrm{E}_{-j} \log p\left(\boldsymbol{v}, \boldsymbol{y} \mid \hat{\boldsymbol{\eta}}\right)\right\}$

  **end**

**until** convergence of all variational parameters;

### S.3 Data-generation design for the simulation studies

Given the remarkably complex and multifaceted biochemical processes involved in genetic regulation, multiple interrelated steps are required to generate realistic datasets with epigenome-induced QTL associations. For each of these steps, we take special care to accommodate a wide range of parameter settings in order to cover a variety of plausible regulatory programs. We will also focus on producing scenarios demonstrating pleiotropy, with hotspots of diverse "sizes" (numbers of associated traits).

The code for generating simulated traits, SNPs (real or simulated) and epigenetic marks (real or simulated), as well as association patterns across these three data types, can be found under the form of documented functions in the R package `echoseq` freely available online[2]; it can be employed to generate alternative association maps to those presented hereafter, under a panel of assumptions from which the user can choose. This resource can also be used as an independent tool to generate epigenome-driven synthetic QTL data which mimic real data conditions.

We base all our numerical experiments on real genetic data which we supplement with data simulated according to generally-accepted principles of population genetics. To fix ideas, we present the steps for the general scenario with $M$ modules; the canonical scenario with no module is readily obtained by choosing $M = 1$.

- **Simulation step 1 — independent loci from real genotyping data**. We start by building the $n \times p$ SNP matrix $\boldsymbol{X}$ by concatenating loci from quality-checked genotyping data for $n = 413$ healthy European individuals with minor allele frequency $> 0.05$[3,4]. Namely, we draw the locus sizes from a Poisson distribution with a prespecified mean, and sample the loci from chromosome one, making sure that they are sufficiently far apart (i.e., separated by at least 150 SNPs which corresponds to a median size of 1 Mb) so as to be reasonably assumed "independent loci".
- **Simulation step 2 — epigenetic control map**. We then form the "control map" between the epigenetic marks and the SNPs. We randomly select up to three active SNPs from each locus, stopping when a prespecified total number of active SNPs has been reached; hence some loci may contain no active SNP. We similarly select $r_0$ active marks among $r$ binary epigenetic marks to be simulated (see below); the remaining $r - r_0$ marks will have no role in the generation of the QTL associations. Not all QTL associations are expected to result from epigenetic modifications. To accommodate this, we specify a proportion of active SNPs whose QTL associations will be triggered by active marks; the remaining active SNPs will have QTL associations simulated independently of the action of the marks. The epigenome control map design is slightly more involved when $M > 1$ modules are simulated, since it must reflect the fact that distinct modules can be governed by distinct epigenetic processes. In other words, the set of active marks and their action on SNP activity are simulated as module-specific, i.e., the active traits within a given module are associated with SNPs whose activity is triggered by specific marks, and these active marks may differ from those triggering QTL associations with traits from another module. The number, $r_{0m}$, of active marks controlling each module corresponds to the minimum between $r_0$ and a draw from a zero-truncated Poisson distribution with parameter 1. All module-specific active marks are then placed randomly among the $r_0$ active marks, ensuring that each of the $r_0$ marks are assigned to at least one module.
- **Simulation step 3 — QTL control map**. We next generate the pleiotropic QTL association pattern. For each active SNP $s$, we choose a subset of active modules (in the non-module case $M = 1$, this step is skipped). Then, for each of these active modules, we draw the proportion of its traits controlled by the SNP from a uniform distribution (first and third simulation study) or

from a right-skewed Beta distribution, favouring large hotspots (second simulation study). We then randomly select the traits associated with SNP $s$ within the module according to this proportion. These module-specific *hotspot propensities* therefore produce hotspots of different sizes within and across the active modules.

- **Simulation step 4 — epigenetic marks**. We then effectively generate a $p \times r$ binary matrix of marks $\boldsymbol{V}$ as follows. For each mark, we draw a proportion of SNPs falling in the mark from a left-skewed Beta distribution (so the mark concerns relatively few SNPs), and we randomly select the SNPs concerned by the mark. We code this mapping in $\boldsymbol{V}$ by assigning the value unity if and only if the SNP (row of $\boldsymbol{V}$) falls within the mark (column of $\boldsymbol{V}$) and we make sure that each mark concerns at least two SNPs. We then enforce that each entry of $\boldsymbol{V}$ corresponding to a pair of active mark and SNP whose activity is triggered by the mark is set to unity.
- **Simulation step 5 — molecular traits**. Given this matrix $\boldsymbol{V}$ and the above epigenome- and QTL-association maps, we generate a series of auxiliary variables in view of simulating the traits as the sum of a genetic component and an independent noise component. For each module $m$, we first obtain an $r \times 1$ regression vector $\boldsymbol{\xi}_m$ for the epigenetic marks, such that its nonzero entries correspond to active marks (i.e., inducing QTL associations with traits from module $m$) and have a log-normal distribution. Hence the marks have non-negative effects, thereby only increasing the potential of SNPs to be involved in QTL activity, but not decreasing it ("enrichment effect"). We then draw $z_{st} \sim \mathcal{N}(\zeta, 1)$, for all $s = 1, \ldots, p$, $t = 1, \ldots, q$, with a large negative mean $\zeta = -2.5$ to induce overall sparsity. For each active SNP $s$ and module $m$ controlled by it, we next proceed as follows: if the QTL activity is triggered by the epigenome, we set $z_{st} \leftarrow z_{st} + \boldsymbol{V}_s^T \boldsymbol{\xi}_m$ for all traits $t \in m$ associated with SNP $s$; if the activity is not triggered by the epigenome, we set $z_{st} \leftarrow z_{st} + \theta_{m,s}$ for all traits $t \in m$ associated with SNP $s$, where $\theta_{m,s}$ is a SNP-specific effect drawn from a log-normal distribution. We then obtain binary variables specifying the QTL association pattern, $\gamma_{st} = \mathbb{1}(z_{st} > q_{1-\alpha})$, for all $s = 1, \ldots, p$, $t = 1, \ldots, q$, where $q_{1-\alpha}$ is the $1-\alpha$ empirical quantile of $z_{st}$ ($s = 1, \ldots, p$, $t = 1, \ldots, q$), with $\alpha$, a chosen proportion of pairwise associations. We next use these variables to generate the regression coefficients $\beta_{st}$. If $\gamma_{st} = 0$, we set $\beta_{st} = 0$. To set the $\beta_{st}$ for which $\gamma_{st} = 1$, we first draw the proportion of a trait's variance explained by individual SNPs from a left-skewed Beta distribution to favour the generation of smaller effects and we then rescale these proportions so that the proportion of genetic variance of each trait does not exceed a prescribed value. The magnitude of the $\beta_{st}$ derives from this value, and its sign is altered with probability 0.5. This implies an inverse relationship between minor allele frequencies and effect sizes, as expected under natural selection[5]. Finally, we build the $n \times q$ matrix of traits, $\boldsymbol{y} \leftarrow \boldsymbol{X}\boldsymbol{\beta} + \boldsymbol{\varepsilon}$, where the noise $\boldsymbol{\varepsilon}$ is a centred multivariate normal variable with covariance such that the traits are equicorrelated with coefficient drawn from the interval $(0, 0.25)$; for module-based scenarios, the noise component is modelled using a multivariate normal variable with block equicorrelation.

Here the number of traits, $q$, is either prespecified or drawn from a Poisson distribution with a given mean; in the third simulation study, we simulate five modules of traits, and the number of traits in each module corresponds to a Poisson draw with mean 50.

We evaluate statistical performance based on 32 data replicates for each scenario, and use the annealing-augmented version of our algorithm, with a geometric schedule for a grid of 100 inverse temperatures and with initial temperature $T = 5$.

### S.4 Addendum to simulation studies: null scenario

We complement the simulation studies with a comparison of EPISPOT and ATLASQTL when none of the 500 candidate marks supplied to the former method contribute to the QTL associations. Table T1 shows the 95% confidence interval of the standardised pAUC for the problems of Section "Performance under varying degrees of epigenome involvement" in the main text, with $p_{\text{epi}} = 0$ (no involvement of the epigenome) and a mean number of traits $\lambda = 200, 400, 600, 800, 1000$ and 1600. No significant difference between the methods is observed as all intervals overlap. This also suggests that supplying many irrelevant marks (noisy information) to EPISPOT does not deteriorate performance compared to methods that do not encode these marks (here ATLASQTL).

| Mean number of traits | ATLASQTL | EPISPOT |
|---|---|---|
| 200 | (0.72, 0.75) | (0.73, 0.76) |
| 400 | (0.74, 0.77) | (0.77, 0.79) |
| 600 | (0.74, 0.78) | (0.76, 0.79) |
| 800 | (0.77, 0.80) | (0.79, 0.82) |
| 1000 | (0.79, 0.83) | (0.82, 0.84) |
| 1600 | (0.80, 0.84) | (0.82, 0.84) |

Table T1: Performance for problems with 500 marks drawn from noise ($p_{\text{epi}} = 0$): 95% confidence interval of the standardised pAUC for ATLASQTL and EPISPOT based on 32 replicates. See the main text for details on the data generation.

### S.5 Details on the monocyte eQTL case study

**S.5.1 Monocyte eQTL datasets, epigenetic annotations and selection of candidate susceptibility loci.** The first dataset consists of genotyping and microarray expression measurements for $n = 413$ healthy European individuals[3,4]. The genotyping data are preprocessed using standard quality control filters (SNPs with call rate $< 95\%$, violating the Hardy–Weinberg equilibrium assumption at nominal $p$-value level $10^{-4}$ and with minor allele frequency $< 5\%$ are discarded). The transcripts are also quality checked and only the levels with mean expression and IQR above the first quartile of their corresponding empirical distribution are kept. This results in retaining $p \approx 550$K SNPs and $q = 22,827$ transcripts for analysis. Known confounders, namely, age, gender and batch, are regressed out from the expression matrix. To account for hidden confounders, we first derive principal components for the transcripts screened as free of any genetic control (i.e., using a preliminary univariate screening with MATRIXEQTL[6] and retaining the transcripts whose association $p$-values with SNPs are all $> 10^{-6}$) and we regress out the first $k = 10$ components from the expression matrix, where $k = 10$ is obtained by maximising the number of *cis* and *trans* associations. Here and throughout the analysis, an effect between a SNP and a transcript is called *cis* eQTL if the SNP and the transcript are on the same chromosome and no more than 1 Mb apart, and it is called *trans* eQTL otherwise.

The application of ATLASQTL at a genome-wide level reveals substantial pleiotropy around the gene *LYZ*, on chromosome 12. To shed light on the genetic mechanisms underlying this pleiotropy, we focus our analysis on the genetic variants located on chromosome 12. We next describe the data preparation steps for the EPISPOT analysis.

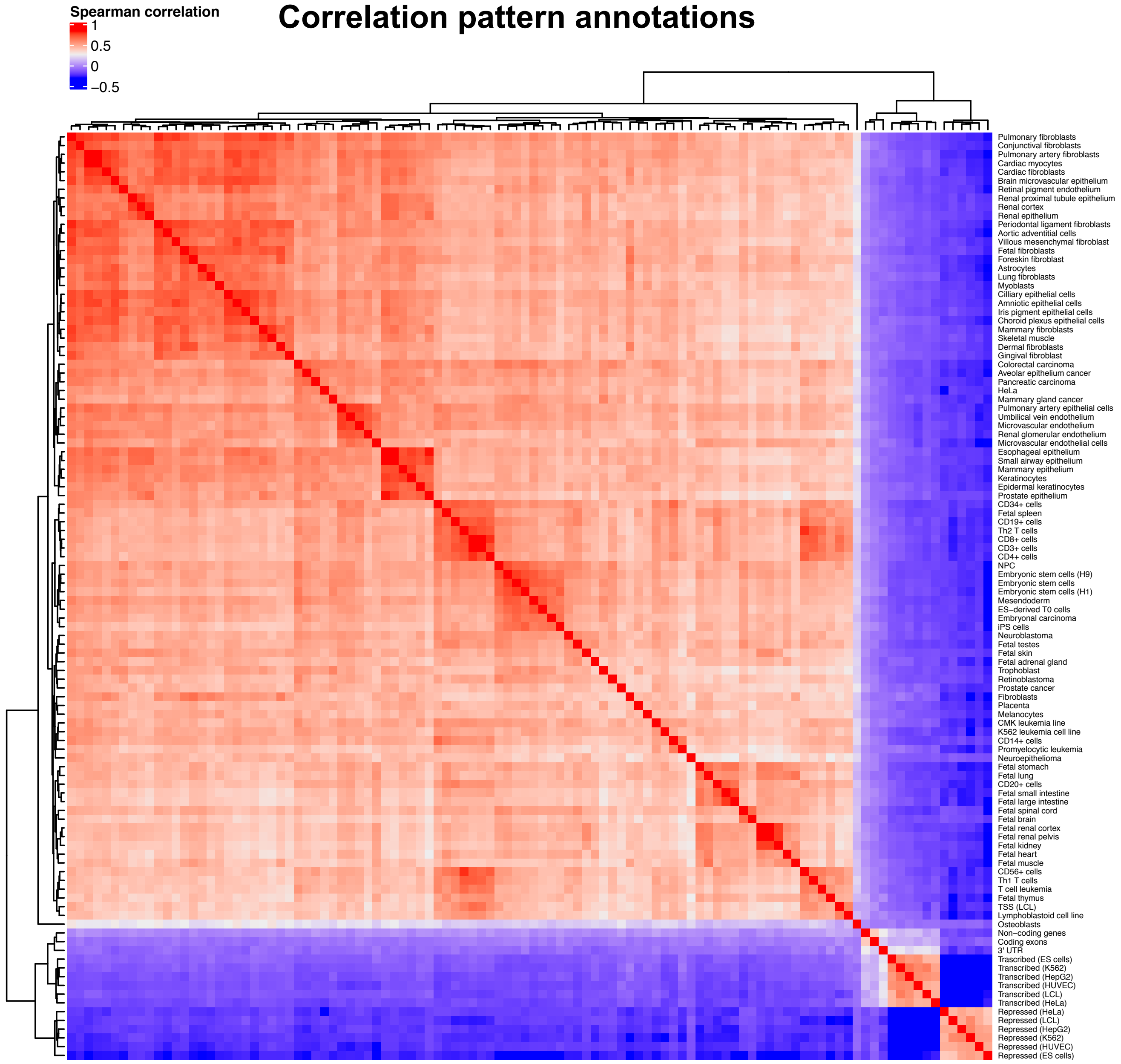


Figure F1: Correlation of the candidate epigenetic annotations supplied to M-EPISPOT. All variables are binary, except the distance to the closest transcription start site (TSS) which is not included in the heatmap.

- **Analysis step 1 — prescreening with ATLASQTL**. We first consider the ATLASQTL run on the first dataset and take note of all QTL associations involving SNPs from chromosome 12 based on a permutation-based Bayesian FDR threshold of 5% (using spline-based interpolation) on the posterior probabilities of inclusion. We obtain 382 *cis*, resp. 595 *trans* associations, involving 350 unique SNPs on chromosome 12 and 430 unique transcripts overall.

Of note, since the ATLASQTL mapping is multivariate, it tends to find sparse association patterns even in case of strong LD, meaning that a large proportion of the 350 identified SNPs is expected to tag for distinct eQTL signals, corresponding to independent active regions. By comparison, the classical univariate mapping approach MATRIXEQTL[6] reported a total of 8 050 eQTL associations, corresponding to 1 054 associated variants at FDR 5%, yet only a fraction of these variants will represent independent signals, as many are in LD and hence likely tagging a same causal SNP.

We also define the *LYZ region* as encompassing all SNPs located $< 1$ Mb upstream or downstream to *LYZ*. SNPs in this region are responsible for 515 *trans* associations and only 22 *cis* associations, reflecting the high level of pleiotropy; in comparison SNPs outside the *LYZ* region control 360 *cis* and 80 *trans* associations.

- **Analysis step 2 — candidate loci**. We then prepare the second, independent monocyte eQTL dataset involving $n = 286$ healthy European individuals from the CEDAR study[7]. These data consist of imputed SNPs (Sanger Imputation Services with the UK10K+ 1000 Genomes Phase 3 Haplotype panels) and $q = 12,771$ quality-checked microarray transcript levels, which are preprocessed using the same filtering- and confounding-adjustment procedure as for the first dataset. We look up all 350 active SNPs in this dataset and we form loci by gathering all imputed SNPs in a 25 Kb neighbourhood of each active SNP and then merging overlapping regions. This results in a total of $p = 1,543$ SNPs distributed into 195 loci on chromosome 12; we concatenate all the loci to form an $n \times p$ matrix $\boldsymbol{X}$ of candidate SNP predictors.

- **Analysis step 3 — epigenetic annotations**. We then retrieve epigenetic information for SNPs from the 1000 genome project, using a curated database[8]. This database gathers different genomic annotations, namely, DNase-I hypersensitivity sites (DHS) for a range of tissues and cell lines, annotations on gene structures ($3'$ and $5'$ UTRs, protein-coding exons), as well as genome segmentation annotations reporting whether nearby histone modifications are in line with transcription start sites (TSSs), CTCF binding sites, enhancer activity, promoter-flanking regions or repressed chromatin. Moreover, the distance of genetic variants to their nearest TSS in the Ensembl gene database is also provided. With the exception of the distance to TSSs, each entry of a given mark is coded unity if the corresponding variant falls in the mark and zero otherwise. We further merge experimental replicates by taking the union of the annotations derived from the same tissue or cell type. We hence obtain a total of $r = 168$ candidate epigenetic annotations for all our candidate SNPs but three, which are excluded from the $\boldsymbol{X}$ matrix and from all downstream analyses. We last drop all binary marks which concern $< 5\%$ of the SNPs in $\boldsymbol{X}$ and gather the remaining marks in a $p \times r$ matrix $\boldsymbol{V}$, where now $p = 1,540$ and $r = 107$; $\boldsymbol{V}$ has no missing entry.

Figure F1 shows the correlation structure among the candidate annotations in $\boldsymbol{V}$. It shows that the annotations tend to group according to their type. The majority of marks pertain to DNase-I hypersensitivity sites (DHS) in different tissues and cell types, and tend to cluster together on the top-left 4/5 of the heatmap. Moreover, DHS in similar tissues/cell types also form subgroups. The remaining marks relate to gene structures and genome segmentation annotations.

The marks selected by M-EPISPOT and their corresponding posterior probabilities of inclusion (epi-PPIs) are provided in Table S2. The relevance of the DHS in $\text{CD14}^+$ monocyte cells and TSS distance annotations is extensively discussed in the main text. The presence of the three marks pertaining to leukemia/K562 cells may result from the propensity of these cells to develop features similar to monocytes[9]. The interpretation of some other marks would deserve further investigation; certain marks may act as proxies for other marks, highly correlated with them.

- **Analysis step 4 — modules of transcripts**. We then look up all 430 active transcripts in the second dataset; 50 transcripts are missing in this dataset and we assign the remaining 380 transcripts to two "modules" based on whether they were controlled by SNPs from the *LYZ* region in the first dataset (*pleiotropic module*, for "pleiotropic" QTL control) or not (*scattered module*, for "scattered" QTL control). We then augment each module by adding all transcripts highly correlated with any transcript in the module, starting with the *pleiotropic module* (Pearson correlation $\rho > 0.9$). This results in a partition with $q = 283 + 191$ transcripts in the *pleiotropic* and *scattered modules*,

| | Mean # assoc. transcripts per SNP | Mean # assoc. transcripts per *active* SNP | Maximum # assoc. transcripts per SNP |
|---|---|---|---|
| $\boldsymbol{Y}$ | 17.61 | 79.25 | 154 |
| $\boldsymbol{R}_{\text{LYZ}}$ | 7.61 | 34.25 | 134 |
| $\boldsymbol{R}_{\text{CREB1}}$ | 2.44 | 14.67 | 36 |

Table T2: Summary of the number of transcripts associated with each SNP from the *LYZ* pleiotropic locus, using a permutation-based FDR of 5%. These numbers are obtained from three M-EPISPOT runs using $\boldsymbol{Y}$, $\boldsymbol{R}_{\text{LYZ}}$ and $\boldsymbol{R}_{\text{CREB1}}$, respectively, as matrix of molecular traits. Column one shows the average numbers across all SNPs while column two only consider the SNPs with at least one association (active SNPs).

| | Nb of edges | Density (%) | Mean degree | Max degree | Modularity |
|---|---|---|---|---|---|
| $\boldsymbol{Y}$ | 2338 | 2.12 | 9.95 | 58 | 0.47 |
| $\boldsymbol{R}_{\text{LYZ}}$ | 1756 | 1.59 | 7.44 | 41 | 0.56 |
| $\boldsymbol{R}_{\text{CREB1}}$ | 706 | 0.64 | 3.00 | 16 | 0.75 |

Table T3: Summary of the transcript conditional independence networks for $\boldsymbol{Y}$, $\boldsymbol{R}_{\text{LYZ}}$ or $\boldsymbol{R}_{\text{CREB1}}$ using the graphical model method `beam`[10].

respectively. We gather all transcripts to form an $n \times q$ matrix $\boldsymbol{y}$ of traits.

- **Analysis step 5 — settings for the QTL methods**. Finally, to ensure common comparative grounds, we use the same settings for all the methods (M-EPISPOT, EPISPOT and ATLASQTL), i.e., annealing schemes with same schedule, as well as a prior average number of SNPs associated with each trait of 2 and a corresponding prior variance of 4 (Section S.1).

We base the replication of the ATLASQTL prescreening results on the transcript levels available in CEDAR and we map the hits up to proxy SNPs in a 1 Mb window.

#### S.5.2 Genetic association and network analyses of the *LYZ* hotspot mediation effects.

In this section, we evaluate plausible mediation mechanisms between the pleiotropic *LYZ* locus (Figure 5D-i, main text) and the controlled transcripts from the so-called *pleiotropic module*, as described in the monocyte eQTL case study. Our analyses have highlighted two candidate gene mediators for the action of the hotspots in this locus, namely, *LYZ* and *CREB1*. We evaluate these hypotheses based on three types of considerations.

First, we assess the presence of a genetic signal after correcting for the effects of each of the *LYZ* and *CREB1* genes from the matrix of expression levels. Table T2 gives a summary of the genetic signals that remain, i.e., by running M-EPISPOT using as matrix of traits the original transcript levels ($\boldsymbol{Y}$), the residual levels after regressing out the effects of the *LYZ* transcripts ($\boldsymbol{R}_{\text{LYZ}}$), and the residual levels after regressing out the effects of the *CREB1* transcripts ($\boldsymbol{R}_{\text{CREB1}}$). It indicates that the hotspot activity is somewhat reduced after removal of the *LYZ* transcript effects, but substantially more after removal of the *CREB1* transcript effects. In this latter case, the top hotspot size has been reduced from 154 in the original M-EPISPOT analysis to 36, using an FDR of 5%. This speaks in favour of a mediation role of *CREB1*, possibly in conjunction with other genes, such as *LYZ*.

Our second investigation uses a network analysis of the analysed transcripts. Here, we compare the conditional independence graph of the original transcript levels $\boldsymbol{Y}$ with those of the residual levels $\boldsymbol{R}_{\text{LYZ}}$ and $\boldsymbol{R}_{\text{CREB1}}$. To this end, we employ the scalable joint Gaussian graphical method `beam`[10], which implements network analyses based closed-form Bayes factors, along with multiplicity-adjusted edge selection. The results are summarised in Table T3 and Figure F2. They indicate

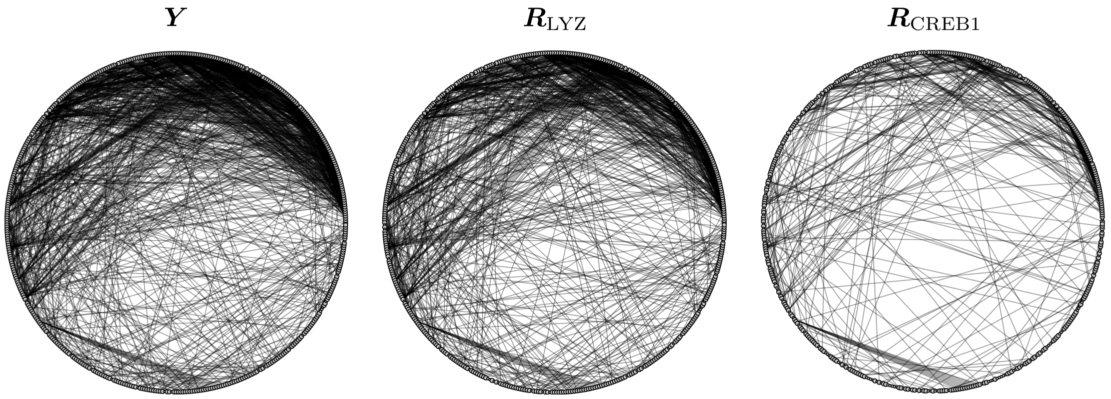


Figure F2: General representation of the transcript conditional independence networks for $\boldsymbol{Y}$, $\boldsymbol{R}_{\mathrm{LYZ}}$ and $\boldsymbol{R}_{\mathrm{CREB1}}$ using the graphical model method `beam`[10]. The connectivity diminishes in $\boldsymbol{R}_{\mathrm{LYZ}}$ compared to the original transcript levels $\boldsymbol{Y}$, and even more so in $\boldsymbol{R}_{\mathrm{CREB1}}$ compared to $\boldsymbol{Y}$.

that the graph $\boldsymbol{R}_{\mathrm{CREB1}}$ is substantially sparser and has higher modularity, i.e., a larger number of disconnected communities, compared to the denser graphs $\boldsymbol{Y}$ and $\boldsymbol{R}_{\mathrm{LYZ}}$, again in line with a strong mediating action of *CREB1*.

Further, more specific considerations are needed to refine and confirm these mediation hypotheses, e.g., via experimental validation.
